# Sex-specific effects of injury and beta-adrenergic activation on metabolic and inflammatory mediators in a murine model of post-traumatic osteoarthritis

**DOI:** 10.1101/2023.08.15.553402

**Authors:** Ravi K. Komaravolu, Padmaja Mehta-D’souza, Taylor Conner, Madeline Allen, Jessica Lumry, Timothy M. Griffin

**Affiliations:** Aging and Metabolism Research Program, Oklahoma Medical Research Foundation, Oklahoma City, OK, USA, 73104.; Department of Health and Exercise Science, University of Oklahoma, Norman, OK, USA, 73019; Oklahoma City VA Health Care System, Oklahoma City, OK, USA, 73104; Oklahoma Center for Geroscience and the Department of Biochemistry and Molecular Biology, University of Oklahoma Health Sciences Center, Oklahoma City, OK, USA, 73104

**Author notes:** CORRESPONDING AUTHOR: Timothy M. Griffin, Ph.D., Associate Member, Aging and Metabolism Research Program, Oklahoma Medical Research Foundation, 825 NE 13^th^ St, Oklahoma City, OK 73104 USA. AUTHOR CONTRIBUTIONS: Concept and design: RKK, TC, PMD, TMG; acquisition, analysis, and interpretation of data: RKK, PMD, TC, MA, JL, TMG; drafting and critical revision of article: RKK, PMD, TC, MA, JL, TMG; final approval of article: RKK, PMD, TC, MA, JL, TMG.

**Keywords:** infrapatellar fat pad, metabolomics, synovial fluid, lipids, sexual dimorphism

## Abstract

Metabolic processes are intricately linked to the resolution of innate inflammation and tissue repair, two critical steps for treating post-traumatic osteoarthritis (PTOA). Here we used the β-adrenergic receptor (βAR) agonist isoproterenol as a tool to perturb intra-articular metabolism 3.5 weeks after applying a non-invasive single-load compression injury to knees of 12-week-old male and female mice. We examined the acute effects of intra-articular treatment with isoproterenol relative to saline on pain behavior, histology, multiplex gene expression, and synovial fluid metabolomics. Injured knees developed PTOA pathology characterized by heterotopic ossification, loss of tibial and femoral articular cartilage, and infrapatellar fat pad (IFP) atrophy and fibrosis. Isoproterenol modestly increased IFP atrophy and fibrosis, and it also caused sexually dimorphic and injury-dependent effects on IFP and synovium gene expression. In injured joints of female mice, isoproterenol suppressed the upregulation of pro-fibrotic genes and downregulated the expression of adipose tissue genes and pro-inflammatory genes (*Adam17*, *Cd14*, *Icam1*, *Csf1r*, and *Casp1*). Injury substantially altered synovial fluid metabolites by increasing amino acids, peptides, sphingolipids, phospholipids, bile acids, and dicarboxylic acids, but these changes were not appreciably altered by isoproterenol. Mechanical allodynia was also not altered by isoproterenol, although isoproterenol downregulated the expression of nociception-associated genes, *Ngf* and *Tacr1,* in injured IFP-synovium of female mice. Overall, these results suggest that βAR activation functions in a sexually dimorphic manner in PTOA joints. The findings support further exploration of therapeutic strategies that target neuro-metabolic signaling pathways for treating PTOA, particularly in women.

## 1 INTRODUCTION

Osteoarthritis (OA) is a multi-faceted disease that poses substantial challenges for developing effective treatments^1^. There is growing consensus that unresolved joint inflammation involving innate immune processes are central factors driving OA progression and clinical symptoms^2,3^. For example, following an initial injury-induced inflammatory response^4,5^, the pathophysiology of post-traumatic OA (PTOA) is characterized by a positive feedback cycle involving chronic joint tissue damage and aberrant remodeling, the generation of endogenous damage-associated molecular pattern molecules, and low-grade activation of cellular and molecular innate immune responses^6,7^. However, traditional anti-inflammatory approaches to disrupt this cycle have not been effective at treating OA. An alternative approach is to target metabolic processes that suppress inflammation and promote tissue repair. Metabolic substrates and signaling pathways are intricately linked to the resolution of innate immune system activation and tissue repair^8^. Thus, our motivation for the current study was to understand how metabolic processes may be therapeutically targeted to reduce PTOA inflammation and pain.

We used the β-adrenergic receptor (βAR) agonist isoproterenol as a tool to perturb intra-articular metabolism and examine its effect on joint inflammation and tissue homeostasis. The endogenous ligands of βARs are norepinephrine and epinephrine. These catecholamines integrate the actions of the sympathetic nervous system in organ systems throughout the body. βARs and catecholamines are present in synovial joint tissues and implicated in joint homeostasis and OA pathophysiology^9–11^.

We focused on intra-articular βAR stimulation for two primary reasons. First, βAR stimulation suppresses classical M1-like pro-inflammatory responses in macrophages and induces an M2-like transcriptome and immunosuppression^12^. Second, βAR stimulation suppresses leptin secretion from adipocytes^13^ and induces adipose tissue lipolysis^14^. These features of βAR stimulation are biologically coupled, as lipolysis increases trafficking of M2-like macrophages to adipose tissue^15^. Thus, acutely stimulating intra-articular adipose tissue lipolysis may help tip the balance to promote the resolution of chronic inflammation during PTOA. However, increasing intra-articular lipolysis may negatively impact joint health by increasing the concentration of pro-inflammatory lipids in synovial fluid and the surrounding tissues, as recently reported by Valdes and colleagues in individuals with OA^16^.

Understanding the balance between these different βAR-dependent actions may provide new insight into the regulation of joint inflammation. Based on the previously reported immunoregulatory actions of norepinephrine in macrophages, we hypothesized that intra-articular βAR stimulation would suppress PTOA-associated inflammation in the infrapatellar fat pad (IFP) and synovium. We tested this hypothesis in male and female mice subjected to a non-invasive knee compression injury resulting in rupture of the anterior cruciate ligament. We intra-articularly administered isoproterenol or saline to contra-lateral knees 3.5 weeks after injury (Figure 1A). We selected this timepoint because it follows a period of dynamic changes in immune cell populations following injury^17^ and may better represent the early chronic phase of unresolved inflammation. We collected tissues 2 days after treatment to focus on the coupling of metabolic and inflammatory changes without also altering PTOA pathology. Our primary outcome was the acute change in gene expression measures of synovial-IFP inflammation and metabolic homeostasis. Secondary outcomes included acute changes in mechanical allodynia, synovial fluid metabolomics, and IFP fibrosis.

**Figure 1.**
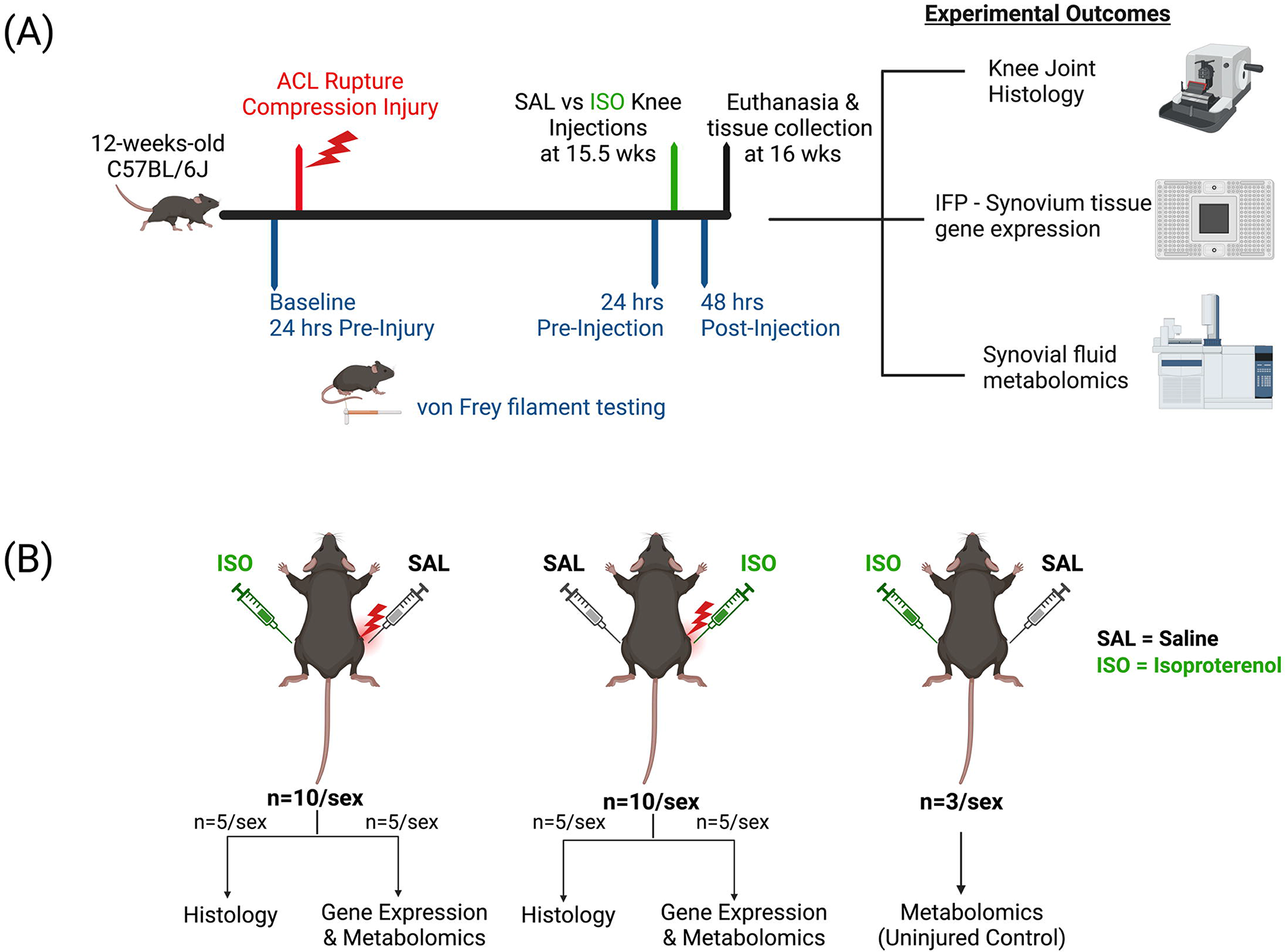
Overview of experimental design. (A) Experimental timeline and outcomes being measured. At 12 weeks of age, 20 male and 20 female C57BL/6J mice were evaluated by von Frey filaments to measure baseline evoked mechanical sensitivity. The next day, mice were anesthetized, and a non-invasive compression injury was applied to the right knee, resulting in rupture of the anterior cruciate ligament. At 15.5 weeks of age, we administered 2µl of sterile saline or filter sterilized isoproterenol hydrochloride (2.5µg/µl; cat. I6504, Sigma-Aldrich) to the injured knee in a randomized, blinded manner, with the contra-lateral knee receiving the opposite solution. We conducted von Frey filament testing 1 day before and 2 days after intra-articular injections. 3 days after injections, animals were euthanized. (B) In half the animals, synovial fluid was collected for metabolomic analysis, and the IFP and adherent synovium was collected for gene expression analysis. In the other animals, limbs were collected for histologic analyses. Three uninjured mice of each sex served as uninjured controls for metabolomic analyses.

## 2 METHODS

### 2.1 Animals and experimental design

Experiments were conducted following protocols approved by AAALAC-accredited Institutional Animal Care and Use Committees at the Oklahoma Medical Research Foundation (OMRF) and Oklahoma City VA. We purchased 10-week-old male (n=23) and female (n=23) C57BL/6J mice from The Jackson Laboratory (Bar Harbor, ME, USA). Animals were group housed at the OMRF vivarium (≤5 animals/cage) in ventilated cages in a temperature-controlled room maintained at 22 ± 3°C on 14h:10h light/dark cycles with *ad libitum* access to chow and water. We habituated animals to the facility for 2 weeks prior to experimentation. An overview of the experimental design is provided in Figure 1, and additional details are provided in Supplemental Materials.

### 2.2 Compression load-induced knee injury

We induced PTOA using a single ramp compression load applied across the knee, resulting in rupture of the anterior cruciate ligament like a previously described non-invasive model of knee PTOA^18^. Mice were anesthetized using isoflurane inhalation while placed in the prone position on a platform adjacent to an electromagnetic materials testing machine (Bose ElectroForce 3100, Eden Prarie, MN, USA) (Figure S1). Prior to loading, animals were administered ketoprofen (5mg/kg, s.c.) for acute pain relief following loading. The right limb was abducted to place the foot in a custom-built holder secured to the bottom platen. The tibia was oriented vertically by placing the flexed knee into a concave holding cup (diameter = 12.3mm; concavity = 2.5mm) secured to the actuated, upper platen. The upper platen was lowered as needed to maintain a 0.6N pre-load for 20 sec. The knee was then compressed by applying a -1.7mm ramp displacement at a rate of -1mm/sec using WinTest Software (TA Instruments, New Castle, DE, USA). Ligament injury was noted in all animals by an audible cue and rapid drop in displacement of the actuated platen (Figure S1). All animals regained ambulatory activity within 30 minutes of the procedure, and no animals required additional analgesic treatment the day after the procedure.

### 2.3 Isoproterenol treatment

3.5 weeks after injury, we performed intra-articular knee injections in isoflurane anesthetized mice. We administered 2µl of sterile saline or filter sterilized isoproterenol hydrochloride (2.5µg/µl; cat. I6504, Sigma-Aldrich) to the injured knee in a randomized, blinded manner, with the contra-lateral knee receiving the opposite solution. Pre-wetted insulin syringes were loaded with solute volumes dispensed onto parafilm using a 0.5-10µl pipette. Based on a synovial fluid volume of 3µl^19^, the effective intra-articular dosing concentration of isoproterenol hydrochloride was estimated to be 4mM. We conducted von Frey filament testing 1 day before and 2 days after intra-articular injections. Three days after injections, animals were euthanized between 9-11AM, and specimens were collected for histology, gene expression, or synovial fluid analyses.

### 2.4 Histological analyses

Left and right knees were isolated as previously described^20^, fixed in fresh 4% paraformaldehyde (pH 7.4) for 24h at 4°C, and decalcified in 10% ethylenediaminetetraacetic acid (EDTA) (pH 7.2-7.4) for 14 days at 4°C. Knees were then dehydrated in an ethanol gradient prior to paraffin embedding and sagittal sectioning. Slides were stained with Hematoxylin, Fast Green and Safranin-O to evaluate structural damage^20^. To evaluate IFP fibrosis, slides were stained in saturated picric acid with 0.1% Sirius red F3B (VWR), washed in 0.5% acetic acid, and counterstained with 0.5% Harris’ hematoxylin (VWR). Two images of the IFP were captured at 4X magnification using a Nikon E200 microscope equipped with a DS-Fi3 digital camera. We used Fiji/ImageJ2 software (v2.9.0/1.54b) to perform RGB color deconvolution and generate a threshold-based binary image from the red channel (e.g. Figure 2B,C). Sirius red staining was quantified as the positive pixel area normalized to the total pixel area of the IFP and adjacent synovium. To evaluate changes in IFP adipose tissue area, we developed a semi-quantitative IFP atrophy score (0-3 scale) as described in Figure S2.

**Figure 2.**
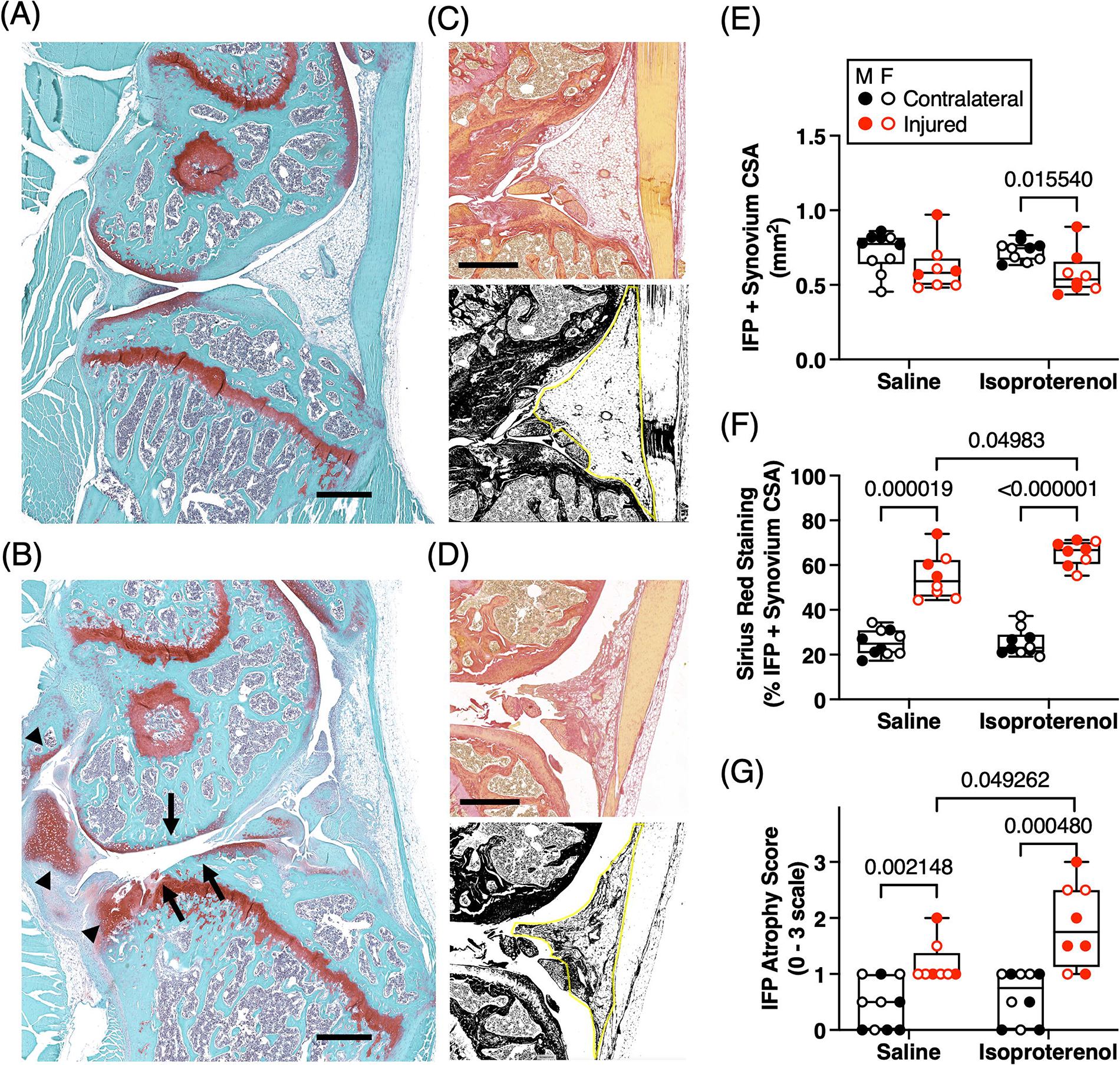
Effect of injury and intra-articular βAR stimulation on knee joint morphology. Representative histological images of knee joints sectioned from an oblique anterior-medial orientation and stained with Hematoxylin/FastGreen/Safranin-O to demonstrate (A) uninjured and (B) injured pathologic changes in the IFP and posterior medial compartment in the same section. Arrows in panel B show cartilage loss to the subchondral bone in the posterior medial compartment, and arrow heads indicate heterotopic chondro-ossifications. Scale bar = 500µm. Representative sagittal sections of (C) uninjured and (D) injured joints stained with Sirius Red to visualize IFP atrophy and fibrosis following injury. Scale bar = 500µm. Bottom panels show corresponding binary images generated from threshold analysis of the red channel following RGB color deconvolution. (E) Cross sectional area (CSA) of the IFP and adherent synovium across all experimental groups, calculated manually as indicated by example yellow lines in binary images from panels C and D. (F) Sirius red staining quantified on binary images as the percent of black pixel area within the total IFP and adjacent synovium CSA. (G) Histologic IFP atrophy and fibrosis score compared across all experimental groups, as described in supplemental Figure S2. Individual animal data (n=5/group per sex) shown for males (closed circles) and females (open circles), except for male/injured/saline and female/injured/isoproterenol (n=3) due to sample preparation issues that prevented usage. Boxes represent the 25th to 75th percentiles, horizontal line indicates the median, and whiskers demonstrate maximum and minimum values. Mann-Whitney or Fisher’s LSD post-hoc paired comparisons shown if p<0.10.

### 2.5 Gene expression and IPA analyses

Immediately following death, the IFP and associated synovium tissue was dissected from each knee, flash-frozen in liquid nitrogen, and stored at -80°C. RNA was isolated from frozen tissue using a TissueLyser (Qiagen), extracted with TRIzol followed by chloroform, and purified using a Clean and Concentrator kit (Zymo Research). 200 ng RNA from each sample was reverse-transcribed into cDNA using an RT2 First-Strand kit (Qiagen). Samples were analyzed by qPCR using a custom DELTAgene Assay that included 90 target genes selected for their role in adipose-tissue homeostasis, fibrosis, inflammation, nociception, and cell adhesion (Table S1). All samples were run on a single microfluidic 96.96 IFC array using a BioMark HD instrument. An isoproterenol treated sample from an injured female animal showed no detection for most genes and was excluded. 5 genes were excluded that did not pass quality control procedures for detection above background in ≥90% of samples. The remaining missing values were imputed using a predicted mean-matching function. Gene expression was quantified relative to the geometric mean of 5 reference (HK) genes (*Actb, Gapdh, Hprt, B2m* and *Rpl14)*.

### 2.6 Synovial fluid metabolomic analyses

Synovial fluid was collected from both knees as previously described^21^ and stored at – 80°C until analysis. Samples were shipped overnight on dry ice to Metabolon (Morrisville, NC, USA) and processed according to their standard pipeline for Global Metabolomic Profiling Analysis (details in Supplemental Methods). The analysis identified a total of 207 named biochemicals within Metabolon’s authenticated standard library (Supplemental Materials), which were detected in nearly all samples (mean=93.6%, median=100%). Peak area data were normalized to extracted volume and then median scaled. Any undetected values (<6.5% of data set) were imputed with sample set minimums on a per biochemical basis.

### 2.7 Mechanical allodynia analyses

To evaluate changes in mechanical allodynia due to injury or βAR stimulation, we conducted von Frey filament testing. Prior to testing, animals were habituated to test chambers for 30 min per day for two consecutive days. On the test day, animals were placed in the chambers for 30 min prior to testing. Seven filaments (sizes: 2.44, 2.83, 3.22, 3.61, 3.84, 4.08, and 4.31) were applied perpendicular to the plantar surface of both hind paws in ascending order, with >1 min between each application. The process was repeated, and the percent positive withdrawal response rate was calculated for each filament and averaged. Mechanical allodynia was indicated by an increase in the overall average percent withdrawal response.

### 2.8 Statistical analyses

Animal group sizes were based on a power analysis for IFP-synovium inflammatory and metabolic gene expression, our primary outcome. Using data from our prior study^22^, n=10 animals per group was estimated to provide 80% power to detect a 1.5-fold change in mean gene expression at p=0.05. The effects of injury and isoproterenol treatment on IFP-synovium histological outcomes gene expression measurements were evaluated by two-way ANOVA. To minimize multiple comparisons, we only performed post-hoc tests to identify between group differences for significant ANOVA analyses (p<0.05). qPCR and synovial fluid metabolomic data were initially assessed by principal component analysis (PCA). We evaluated the loading matrix and partial contribution of variables to identify the primary genes and metabolites, respectively, that contributed to the first principal component. We also performed unsupervised 2-way hierarchical clustering analyses to identify experimental groups that shared similar values of gene expression and metabolite measurements. Expression data for each gene or metabolite were standardized by subtracting the mean and dividing by the standard deviation, and Ward’s minimum variance method was used to calculate cluster distances. Sex-dependent treatment outcomes were evaluated by 2-factor ANOVA and hierarchical clustering analysis under injured conditions. Two-sample comparisons utilized 2-tailed Student’s t-test (p<0.05) unless otherwise noted. Data that did not meet test assumptions for homoscedasticity, even after log transformation, were analyzed by Kruskal-Wallis or Mann-Whitney tests. Group values reported as mean ± S.D. unless otherwise noted. All statistical tests were performed in Prism 9.3.1 (GraphPad Software, Inc) or JMP Pro 16.0.0 (SAS Institute, Inc).

## 3 RESULTS

### 3.1 Knee compression injury induced IFP atrophy and fibrosis

Knee loading injury parameters were similar in male and female animals, including the vertical displacement to rupture (males: -1.370mm ±0.146; females: -1.319mm ±0.147; p=0.2771) and peak compressive force (males: -10.45N ±1.37; females: -9.50N ±1.64; p=0.0547) (Figure S1). Four weeks after loading, male and female mice developed substantial PTOA pathology characterized by heterotopic ossification, loss of tibial and femoral articular cartilage to the subchondral bone in the posterior medial compartment, and IFP atrophy and fibrosis (Figure 2A,B). Given the role of isoproterenol in adipose tissue lipolysis and tissue fibrosis, we investigated the effect of acute intra-articular treatment with isoproterenol (3 days prior to tissue collection) on IFP area, fibrosis, and atrophy in injured and contralateral knees (Figure 2C-G). The cross-sectional area of the IFP and adjacent synovium was reduced with injury by -0.130mm^2^ [-0.0430 to -0.217] (mean difference [95% CI of the difference], p=0.0049), with the strongest effect observed under isoproterenol conditions (Figure 2E). Conversely, Sirius red staining, a marker of tissue fibrosis, was significantly increased in IFP-synovium tissue of injured versus uninjured contralateral knees by 34.79% [30.04 to 39.54] (p<0.0001) (Figure 2F). Moreover, isoproterenol treatment increased Sirius red staining by 5.09% [0.343 to 9.84] (p=0.0364), particularly under injured conditions (injury × treatment interaction: p=0.0360). Injury also caused the IFP to atrophy, which isoproterenol modestly enhanced in the injured knee (p=0.04926) (Figure 2G). We did not observe any sex-dependent differences in these structural outcomes.

### 3.2 Effect of injury and isoproterenol treatment on IFP-synovium gene expression

We performed a PCA using the delta-Ct values from all 85 target genes of all samples to visualize the relative contribution of injury, isoproterenol treatment, and biological sex on variance in IFP-synovium gene expression (Figure 3A). PC1 contributed to 35.5% of the variability in IFP-synovium gene expression, which largely corresponded to injury status. We next performed an unsupervised 2-way hierarchical clustering analysis to compare the expression patterns of the top 20 genes that contributed to PC1 variation across all the samples (Figure 3B). Sample clusters segregated almost completely by injury status, with 16 of the 20 genes significantly downregulated with injury (Figure 3B). Notably, 15 of these 16 downregulated genes are associated with adipose tissue homeostasis, consistent with the observed IFP atrophy following injury. Four genes were upregulated with injury; these genes are functionally related to extracellular matrix proteolysis and cell adhesion.

**Figure 3.**
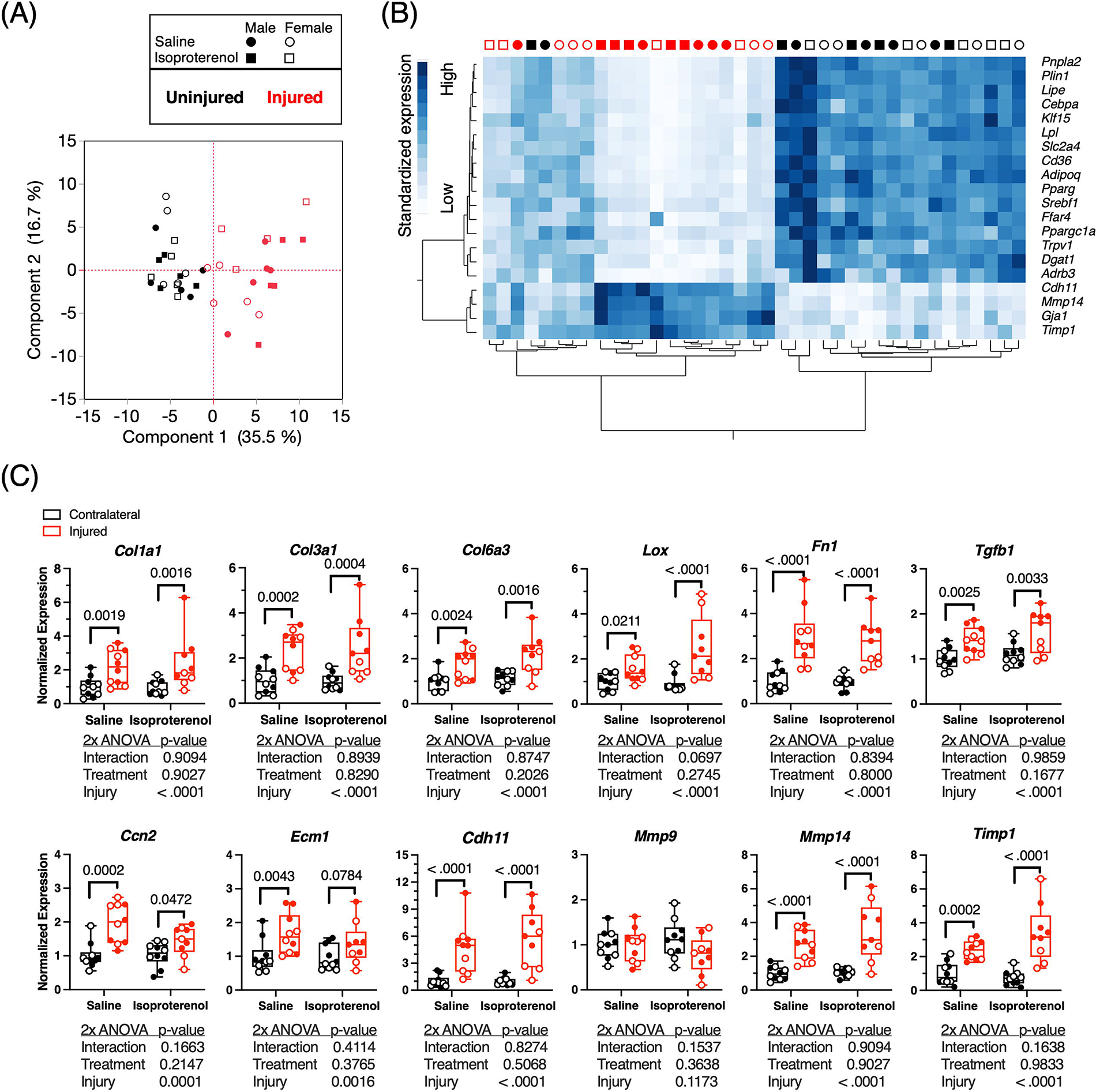
Effect of injury and intra-articular βAR stimulation on IFP-synovium gene expression. (A) Principal component analysis based on delta-Ct data of 85 target genes involved in adipose-tissue homeostasis, fibrosis, inflammation, nociception, and cell adhesion from all experimental groups. Symbols represent individual samples according to treatment (saline=circle, isoproterenol=square), biological sex (male=closed, female=open), and injury status (uninjured=black, injured=red). Principal component 1 (PC1) primarily differentiates between samples isolated from uninjured and injured joints. (B) 2-way hierarchical clustering analysis to compare the expression patterns of the top 20 genes that contributed to PC1 variation across all the samples. Columns represent individual samples according to symbol legend shown in panel A. Heatmap legend signifies standardized gene expression values calculated by subtracting the mean and dividing by the standard deviation. (C) Expression of genes that contribute to connective tissue homeostasis and fibrosis grouped by injury and isoproterenol treatment status. Data normalized to the average expression of contralateral (i.e., uninjured) saline treated samples. Individual animal data shown as closed (male) and open (female) circles. Boxes represent the 25th to 75th percentiles, horizontal line indicates the median, and whiskers demonstrate maximum and minimum values. Mann-Whitney or Fisher’s LSD post-hoc paired comparisons shown if p<0.10 (n=5 per sex per group, except n=4 for female/injured/isoproterenol).

Given the effect of injury and isoproterenol on histologic measurements of IFP fibrosis, we next evaluated these effects on the expression of fibrosis-related genes (Figure 3C). Injury significantly increased the expression of 11 out of 12 genes (*Col1a1, Col3a1, Col6a3, Lox, Fn1, Tgfb1, Ccn2, Ecm1, Cdh11, Mmp14,* and *Timp1*). Although we did not observe a significant effect of isoproterenol treatment on any of the fibrosis-related genes, a visual inspection of the male and female expression patterns (closed versus open circles, respectively) suggested that the effect of injury was suppressed in female mice treated with isoproterenol (Figure 3C). Indeed, when we analyzed the effects of injury and sex by 2-factor ANOVA separately for saline and isoproterenol conditions, we observed significant injury-sex interactions (p<0.05) for half of the genes (*Col1a1, Col3a1, Fn1, Tgfb1, Ccn2*, and *Timp1*) only under isoproterenol conditions. Thus, these findings indicate that isoproterenol suppressed the effect of injury on the expression of fibrosis-related genes in female but not male mice (Figure S3).

### 3.3 Sex-dependent effects of isoproterenol on adipose tissue homeostasis and pro-inflammatory gene expression in injured IFP-synovium tissues

We next analyzed the sex-dependent effects of isoproterenol treatment on the expression of genes involved in adipose tissue homeostasis and inflammation from injured IFP-synovium (Figure 4A,B). Compared to saline, isoproterenol treatment reduced the expression of 12 out of 14 adipose tissue associated genes, including *Adipoq, Adrb3, Cd36, Cebpa, Foxo1, Lep, Lipe, Lpl, Plin1, Pnpla2, Slc2a4, and Zfp423*. Post-hoc analyses showed that the inhibitory effect of isoproterenol on adipose gene expression primarily occurred in female mice (Figure 4A). Notably, female mice expressed significantly higher levels of *Adrb3*, the gene encoding the βAR in adipose tissue, compared to males (sex effect p-value=0.0029; Figure S4).

**Figure 4.**
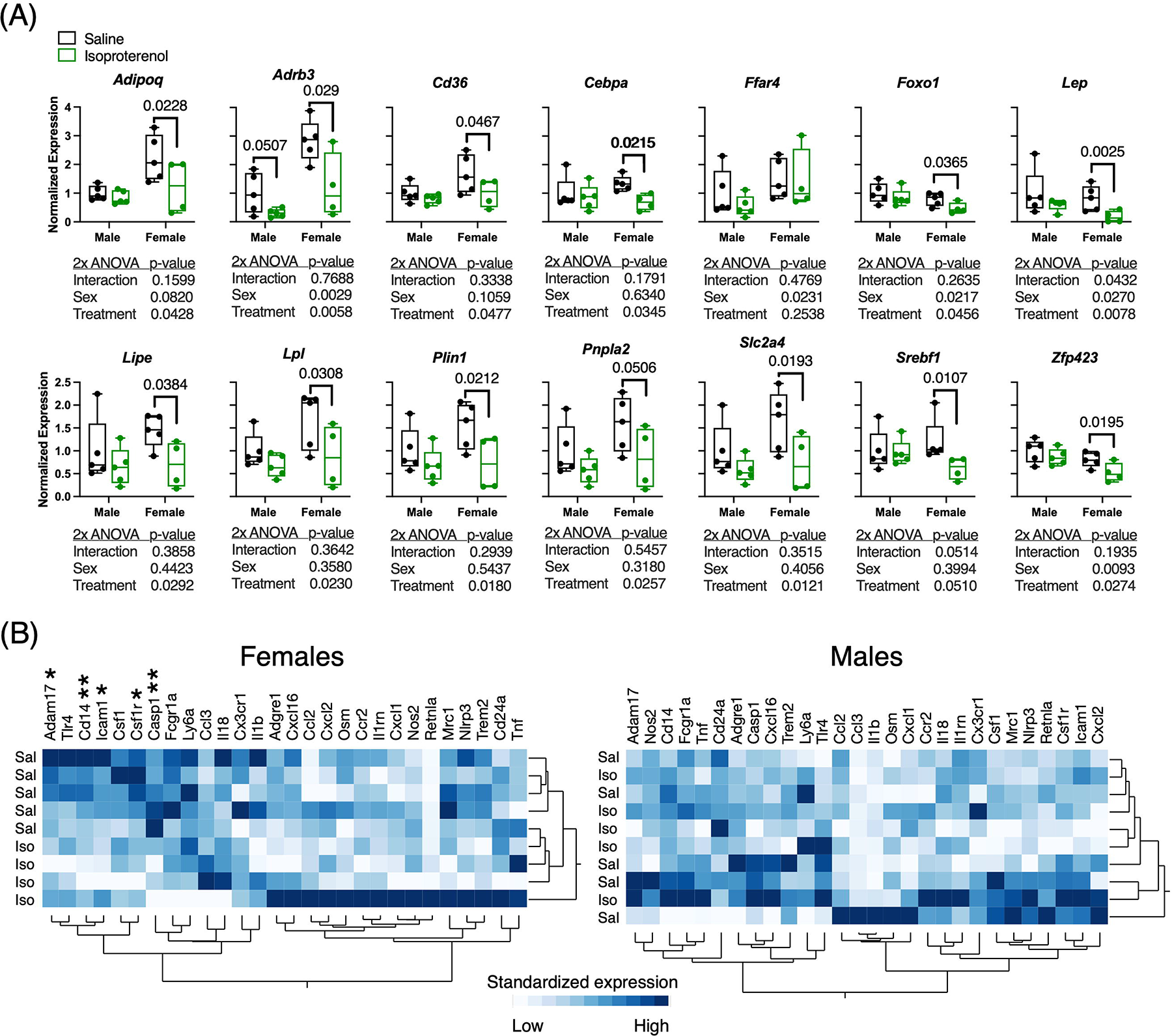
Effect of intra-articular βAR stimulation on the expression of adipose homeostasis and inflammation genes in injured IFP-synovium samples of male and female mice. (A) Expression of genes that contribute to adipose tissue homeostasis and fibrosis grouped by sex and isoproterenol treatment status. Data normalized to the average expression of saline treated samples for male mice. Note that data only include injured IFP-synovium samples. Individual animal data shown as closed circles. Boxes represent the 25th to 75th percentiles, horizontal line indicates the median, and whiskers demonstrate maximum and minimum values. Mann-Whitney or Fisher’s LSD post-hoc paired comparisons shown if p<0.10 (n=5 per sex per group, except n=4 for female/isoproterenol). (B) 2-way hierarchical clustering analysis to compare the expression patterns of immuno-modulatory target genes across saline and isoproterenol treated samples. Saline and isoproterenol samples cluster separately in female but not male mice, and isoproterenol treatment reduced the expression of several genes in female samples as indicated by asterisks (Student’s t test, *p<0.05, **p<0.01). Heatmap colors represent standardized gene expression values as indicated in Figure 3B.

To explore potential sex-dependent effects of isoproterenol on inflammation, we performed unsupervised 2-way hierarchical clustering analyses separately in male and female samples focusing on inflammation-associated gene expression (Figure 4B). Unlike male samples, female samples clustered by saline versus isoproterenol treatment. This clustering pattern showed that isoproterenol reduced the expression of a subset of the inflammatory genes in female mice relative to saline, which was confirmed by Student’s t-test for *Adam17* (p=0.0342), *Cd14* (p=0.0093), *Icam1* (p=0.0361), *Csf1r* (p=0.0196), and *Casp1* (p=0.0032).

### 3.4 Effect of compression injury on global changes in synovial fluid metabolites

We performed a PCA using the normalized peak area data from all detected biochemicals to visualize the relative contribution of injury, isoproterenol treatment, and biological sex on variance in synovial fluid metabolites (Figure 5A). PC1 contributed to 54.2% of the variability in synovial fluid metabolomics, which strongly corresponded to injury status. The metabolites that contributed to PC1 were color-coded by their metabolic super pathway designation and plotted in a descending bar graph based on the percent contribution of each metabolite to PC1 (Figure 5B). This analysis showed that amino acids and peptides were the primary classes of metabolites that contributed to variation in PC1 and thus the effect of injury. When we specifically evaluated the effect of injury by Student’s t-test, irrespective of isoproterenol treatment or sex, we identified 135 upregulated metabolites (q^FDR^<0.01) and 30 downregulated metabolites (q^FDR^<0.003) (Figure 5C, Table S2). This analysis confirmed that injury significantly altered the abundance of many amino acids and peptides, and it further showed that most of these metabolites were upregulated in synovial fluid 4 weeks following injury. The analysis also indicated that synovial fluid lipids showed both upregulation and downregulation following injury. Given the important role of lipids in modulating inflammation, we next focused on how specific lipid sub classes were altered with injury.

**Figure 5.**
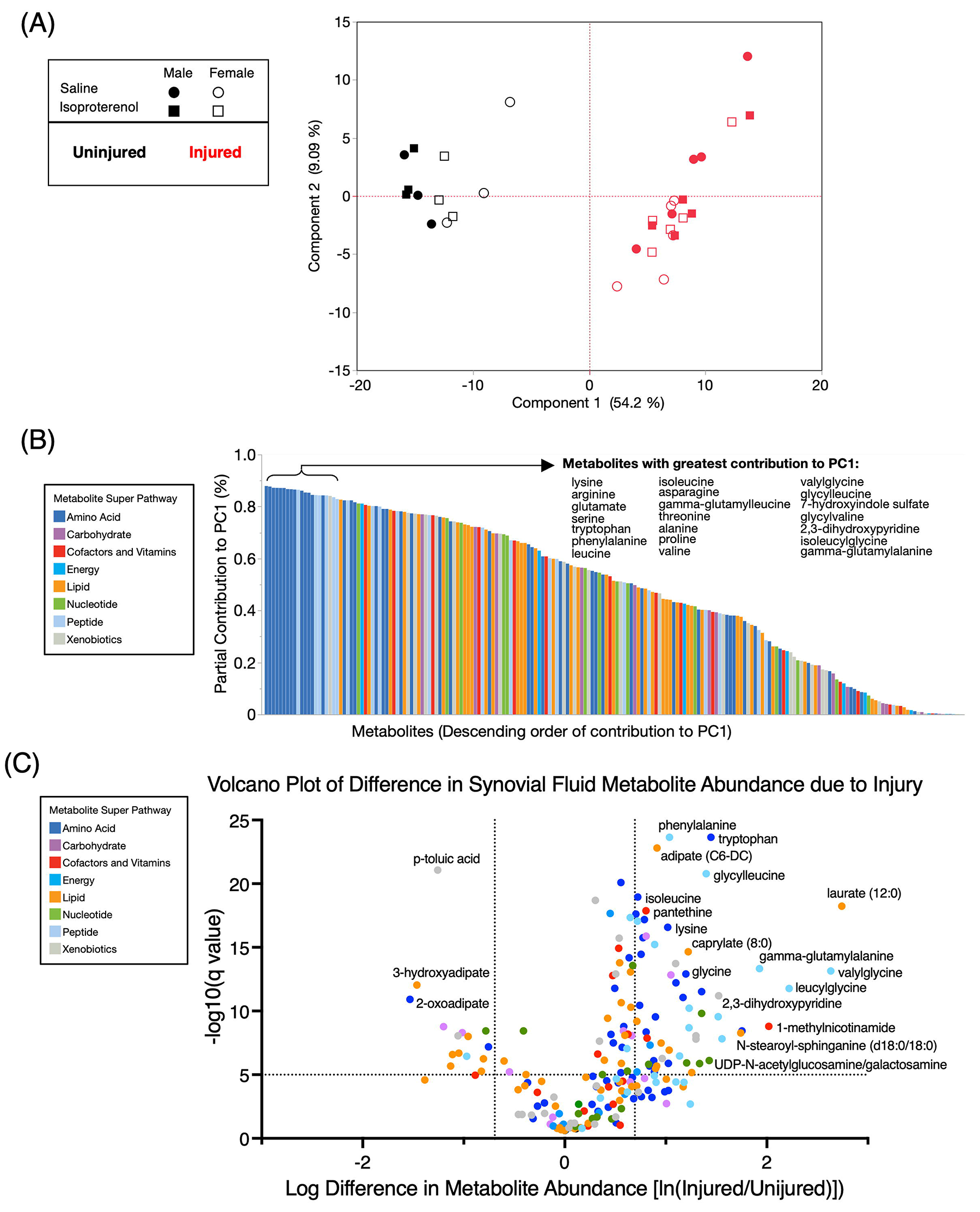
Effect of injury on synovial fluid metabolites. (A) Principal component analysis based on area-under-the-curve relative quantification of 207 metabolites identified in the synovial fluid from a Global Metabolomic Profiling Analysis (Metabolon). Symbols represent individual samples according to treatment (saline=circle, isoproterenol=square), biological sex (male=closed, female=open), and injury status (uninjured=black, injured=red). Principal component 1 (PC1) explained 54.2% of the variance and primarily differentiated between samples obtained from uninjured and injured joints. (B) Bar plot of all identified metabolites ordered in descending contribution to PC1 from left to right. Individual metabolites were colored by their Metabolic Super Pathway assignment (Metabolon), with the top 21 contributing metabolites (primarily amino acids and peptides) listed above the plot in descending text columns from left to right. (C) Volcano plot showing the statistical probability of the log difference in synovial fluid metabolite abundance between injured versus uninjured limbs. Y-axis shows the -log10 false discovery rate (1%) adjusted q-value based on the two-stage step-up method of Benjamini, Krieger, and Yekutieli (dashed line set at q=0.00001). X-axis shows the log difference in metabolite abundance. The vertical dashed lines set at ±0.69314 correspond to a ±2.0-fold difference in abundance.

### 3.5 Effect of compression injury on synovial fluid lipids

We identified 48 synovial fluid biochemicals belonging to the “lipid” super pathway of metabolites. Using the normalized peak area data for these lipids, we performed an unsupervised 2-way hierarchical clustering analysis across all the samples (Figure 6). The analysis generated two major sample clusters, which separated based on injury status. Sex-specific sub-clusters were observed within uninjured, but not injured, samples. Isoproterenol-dependent clustering was not observed. Across the 48 lipids, we identified three clusters based on whether they increased with injury (27 lipids), decreased with injury (15 lipids), or were not altered with injury (6 lipids). Lipids that were more abundant in the synovial fluid of injured knees included straight chain fatty acids, sphingolipids, phospholipids, bile acids, medium and long-chain dicarboxylic acids, and 2 dicarboxylic/hydroxy fatty acids. The lipids with the greatest fold-change increase included the straight chain fatty acid laurate (12:0) (15.5-fold [13.7–17.3]) (mean [95% CI]) and the sphingolipid N-stearoyl-sphinganine (d18:0/18:0) (5.05-fold [4.39-5.71]. In contrast, lipids that were less abundant in injured knees included several long-chain diacylglycerols, short-chain dicarboxylic acids, hydroxy fatty acids, and 4 dicarboxylic/hydroxy fatty acid species. The lipids with the greatest fold-change decrease were the dicarboxylic/hydroxy fatty acids 13-HODE + 9-HODE (0.19-fold [0.15-0.22]) and 3-hydroxyadipate (0.23-fold [0.20-0.26]).

**Figure 6.**
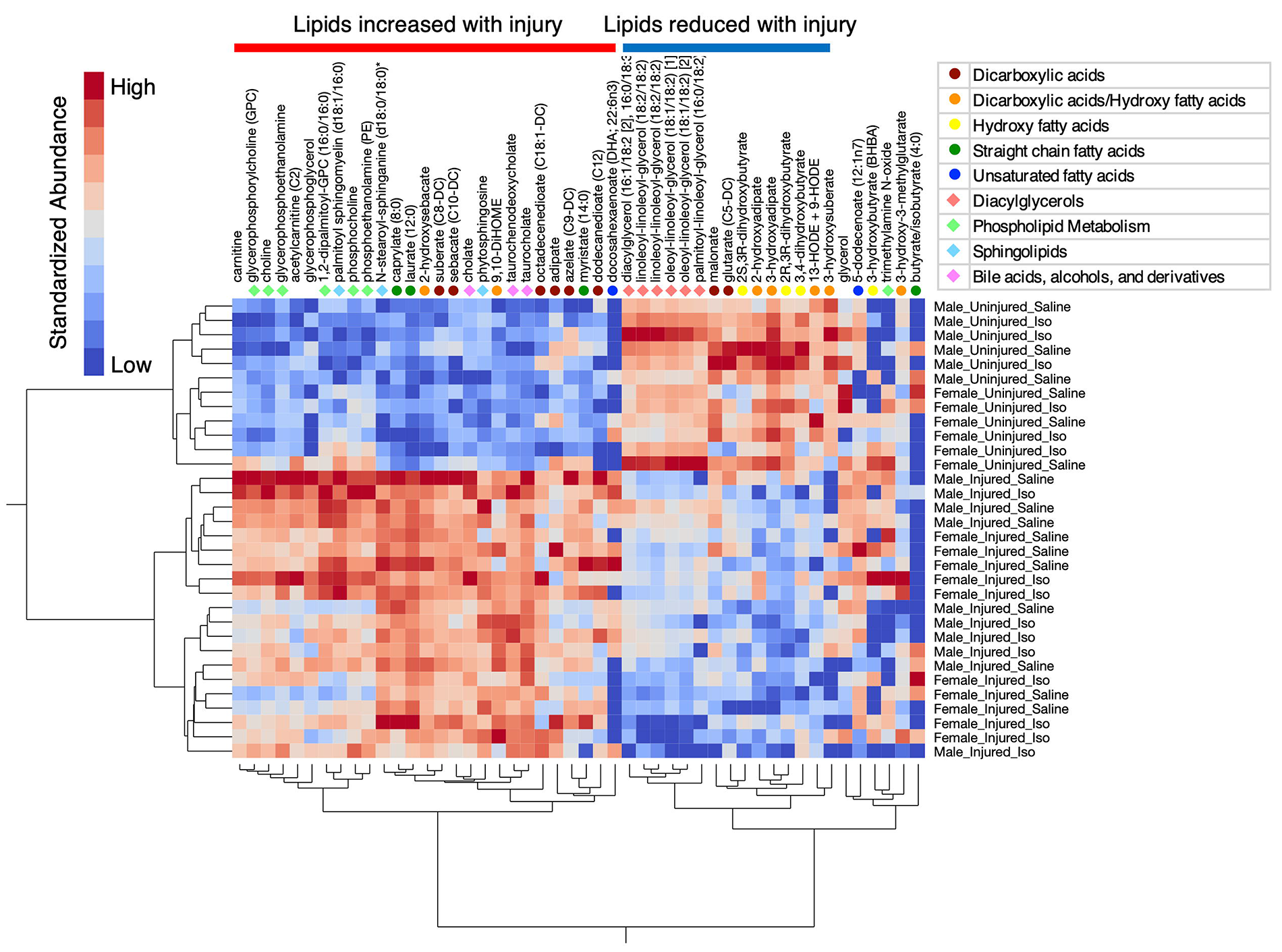
Injury differentially alters the abundance of specific classes of lipids in synovial fluid. 2-way hierarchical clustering analysis was used to identify pattens among the experimental groups according to the relative abundance of synovial fluid metabolites assigned to the lipid super pathway (Metabolon). Columns represent individual metabolites, with lipid sub-pathway labels added as indicated in the legend. Rows represent individual samples labeled by experimental group, which clustered into samples obtained from uninjured and injured knees. The analysis further showed that 27 lipids in the left metabolite cluster increased with injury (p<0.05 confirmed by Student’s t test). 15 lipids in the right metabolite cluster decreased with injury (p<0.05), and 6 lipids did not change with injury. Heatmap color legend signifies standardized metabolite abundance values calculated by subtracting the mean and dividing by the standard deviation.

### 3.6 Sex-dependent effects of isoproterenol on mechanical allodynia and expression of nociception genes

Prior to injury, the average von Frey filament withdrawal response rate was 40.5%±3.3 for males and 48.4%±0.8 for females. 3.5 weeks after injury, the withdrawal response rate substantially increased to 63.7%±5.3 for males and 70.4%±6.6 for females (average of left and right limbs). The withdrawal response of the injured limb compared to the uninjured contralateral limb was significantly greater in female mice (p=0.0145) and showed a similar trend in male mice (p=0.0621) (Figure 7A,B). We evaluated the effect of isoproterenol administration relative to saline 48 hours after intra-articular injection in both injured and contralateral limbs. Neither male nor female mice showed a change in withdrawal response in the injured or contralateral limb due to isoproterenol treatment compared to saline (Figure 7A,B). However, the percent change in withdrawal responses was increased in both limbs of female, but not male, mice following saline and isoproterenol injections (p<0.05), suggesting an overall increase in sensitization in female mice. To further evaluate sex-specific effects of isoproterenol on nociception, we measured the expression of two genes involved in regulating nociception, *Ngf* and *Tacr1*, in IFP-synovium tissue. Both genes were expressed less in female versus male mice (Figure 7C), and there was a trend for a sex × treatment interaction for both genes (p<0.10). Post-hoc analysis indicated that isoproterenol treatment further downregulated *Ngf* and *Tacr1* only in female mice.

**Figure 7.**
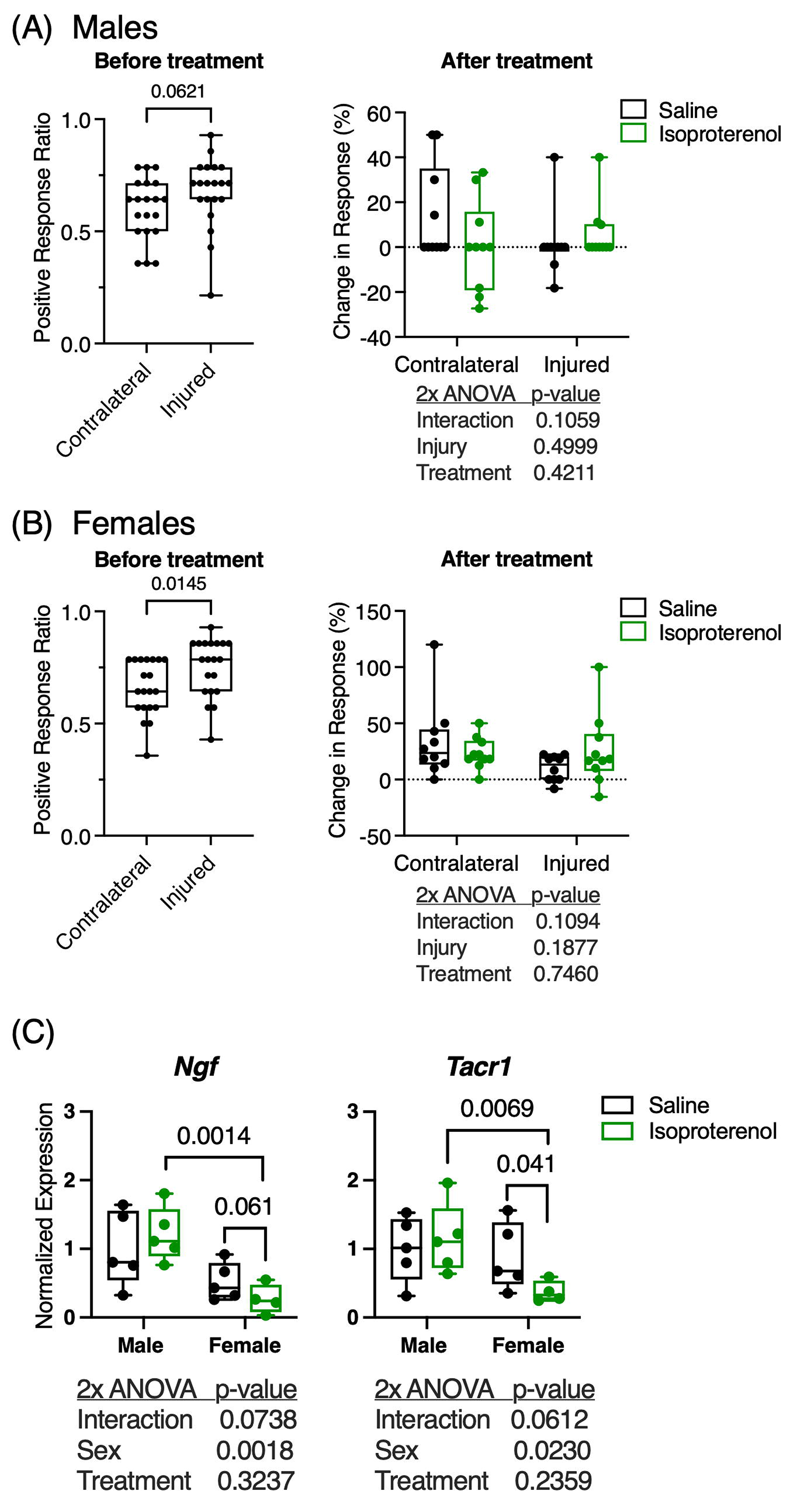
Effect of acute intra-articular βAR stimulation on mechanical allodynia and nociception-related gene expression. Box plots of the ratio of positive withdrawal responses compared to all filaments tested in (A) male and (B) female mice. Data collected 3.5 weeks after injury and 1 day prior to intra-articular isoproterenol injection (symbols represent individual animals, n=10/group per sex, p-values from Mann-Whitney test). 48 hours after isoproterenol treatment, the percent change in average withdrawal response was calculated based on pre-treatment values and plotted by injury and isoproterenol treatment status for (A) males and (B) females. Boxes represent the 25th to 75th percentiles, horizontal line indicates the median, and whiskers demonstrate maximum and minimum values. (C) Box plots of the expression of genes that encode proteins involved in nociception grouped by sex and isoproterenol treatment status. Data normalized to the average expression of saline treated samples for male mice and only include injured IFP-synovium samples. Fisher’s LSD post-hoc paired comparisons shown if p<0.10 (n=5 per sex per group, except n=4 for female/injured/isoproterenol).

## 4 DISCUSSION

Numerous aspects of metabolism regulate inflammatory processes^23,24^, and metabolic substrates and signaling pathways are intricately linked to the resolution of innate inflammation and tissue repair^8^. The goal of this study was to investigate metabolic processes that may be therapeutically targeted to reduce PTOA inflammation and pain. We evaluated the effect of the βAR agonist isoproterenol because it induces adipose tissue lipolysis and stimulates an M2-like immunoregulatory phenotype in macrophages^12,14^. We hypothesized that intra-articular isoproterenol treatment would suppress PTOA-associated inflammation in the IFP and synovium, and we tested this hypothesis in male and female mice subjected to a non-invasive knee compression injury. Compared to saline, intra-articular injection of isoproterenol reduced the expression of several pro-inflammatory genes (i.e., *Adam17*, *Casp1*, *Cd14*, *Csf1r*, and *Icam1*) in IFP-synovium tissue harvested from knees of female mice with PTOA. Isoproterenol also reduced the expression of adipose homeostasis genes and suppressed the expression of pro-fibrotic genes in IFP-synovium tissue of female mice with PTOA. We did not observe effects of isoproterenol in uninjured knee joints of male or female mice. Overall, these results support our hypothesis and suggest that βAR activation functions in a sexually dimorphic manner in PTOA joints.

Metabolic links to OA involve changes in cellular metabolism, synovial fluid metabolites, and paracrine metabolic signaling from systemic and intra-articular adipose tissue^25–27^. From this perspective, the IFP is a particularly interesting metabolic joint tissue as it is both a site of joint inflammation and a paracrine source of adipokines and synovial fluid metabolites, especially lipids^28,29^. It was recently reported that individuals with greater IFP volume one year after anterior cruciate ligament reconstruction surgery are more likely to develop patellofemoral bone marrow lesions and tibiofemoral cartilage lesions at 1- and 5-years post-surgery, respectively^30^. Thus, treatments that stimulate IFP lipolysis may be beneficial. However, other studies have failed to show an association between IFP volume and knee OA pain^31^, and some studies report that reduced IFP size and signal intensity alteration predict knee OA progression and total knee replacement^32–34^. The lack of consensus regarding IFP size and risk of OA progression may be due to several confounding factors, such as a smaller IFP volume in women than men even after body mass index or body weight normalization^35^, different measurement methodologies^36^, or OA endotype specificity (e.g., post-traumatic, obesity, aging)^37^. IFP size alone may not capture important structural, cellular, and biochemical factors that contribute to OA pathology. Moreover, as suggested by our findings, sex-specific analyses could be important for studies investigating immuno-metabolic mechanisms linking IFP structure, synovitis, and OA.

While previous studies have highlighted sex-dependent mechanisms linking metabolic factors, such as leptin or a high-fat diet, to OA pathology^27,38,39^, our current study suggests a sex-dependent role for sympathetic nervous system signaling. βARs and catecholamines are detected in tissues throughout the joint and alter cellular function *in vitro*^9,10^, implicating a role in joint tissue homeostasis. Indeed, chemical ablation of peripheral sympathetic nerves in male mice subjected to destabilization of the medial meniscus confirmed the contribution of the sympathetic nervous system to PTOA^11^. Suppressing peripheral sympathetic nerve activity exacerbated cartilage calcification and subchondral bone thickening but did not alter cartilage degradation or synovitis, signifying that βAR activation may be a therapeutic strategy for PTOA^11^. The study did not evaluate the IFP, and since it was only conducted in male mice, it is not clear if the results apply to females.

We observed that expression of the primary adipocyte βAR, *Adrb3*, was greater in female IFP-synovium samples compared to males under all experimental conditions. This sex-specific difference in βAR expression may contribute to the sexually dimorphic response to isoproterenol characterized by female sensitivity and male resistance. However, βAR expression does not fully explain our findings because the effects of isoproterenol were primarily observed in injured knees, and injury downregulated the expression of *Adrb3* in male and female samples (Figure S4). Interestingly, isoproterenol treatment further downregulated *Adrb3* only under injury conditions. Inflammation and βAR ligand exposure contribute to adipocyte catecholamine resistance with obesity^40^. However, in the IFP, injury appears to induce catecholamine sensitivity despite down-regulation of the receptor. Recent studies describe how neuron-associated macrophages in adipose tissue regulate lipolysis and adipose tissue homeostasis by removing and degrading interstitial norepinephrine^41,42^. Macrophages are important cellular mediators of PTOA pathology^7^, but their contribution to intra-articular catecholamine signaling and IFP lipolysis has not yet been studied. This emerging area of investigation, termed “neuroimmunometabolism”^43^, may be particularly relevant to PTOA pathology.

A useful approach for investigating changes in joint metabolism during the development of OA is to measure the abundance of synovial fluid metabolites. For example, a targeted analysis of oxylipins in synovial fluid from individuals with and without knee OA showed that two oxylipin products of arachidonic acid, 11,12-dihydroxyeicosatrienoic acid (DHET) and 14,15-DHET, are positively associated with prevalent knee OA and structural OA progression^16^. This finding is consistent with clinical and animal model studies that indicate arachidonic acid is elevated in OA IFP samples^44,45^, suggesting a potential link between IFP lipolysis and synovial fluid metabolites. Clinical analyses have also shown that numerous phospholipid and sphingolipid species are elevated in the synovial fluid of individuals with knee OA. However, the clinical significance of this finding is not well understood due to the various contributions of these lipids to joint lubrication, redox scavenging, and inflammation^46^. Given that we also observed greater phospholipid and sphingolipid metabolites in the synovial fluid of injured versus uninjured mouse knees, this animal model may be useful for investigating the functional contribution of specific phospholipid and sphingolipid species to PTOA. Future work is likewise needed to understand the functional consequences of a reduction in numerous diacylglycerol and hydroxy fatty acid species in the synovial fluid of injured versus uninjured knees.

In addition to lipids, many other metabolites are differentially associated with joint trauma and OA severity^21,47,48^, indicating that broad changes in joint tissue metabolism accompany OA progression. Our metabolomic analysis revealed that the greatest effect of injury was to increase the abundance of synovial fluid amino acids and peptides, including branched chain amino acids. This finding is of particular interest because branched chain amino acids are ligands for the mechanistic target of rapamycin complex 1 (mTORC1) signaling pathway, which promotes the development of PTOA^49^. Although injury profoundly altered metabolite concentrations in synovial fluid, few differences were observed between isoproterenol and saline treatments or between males and females (Figures S5 and S6, respectively). Thus, the sexually dimorphic responses to isoproterenol were not likely mediated by differences in synovial fluid metabolites.

Our study identified new evidence regarding the role of biological sex and βAR stimulation in joint homeostasis using a non-invasive compression injury model in which male and female mice consistently develop PTOA. Despite these advances, there are several limitations. We evaluated a single timepoint in a single model of PTOA, and therefore the effects of βAR stimulation could be different at earlier or later timepoints or in a different model. Repeated treatments and longer follow up durations would also be needed to determine effects on PTOA progression. In addition, further research is needed to conduct a broader analysis of how intra-articular βAR stimulation affects cellular phenotypes and tissues besides the IFP. Finally, although we identified numerous differences between males and females, the current study was not statistically powered to evaluate sexually dimorphic responses. Therefore, it is possible that other biologically meaningful sex differences were not detected.

Despite these limitations, this study provides important new perspectives on how metabolic processes may be therapeutically targeted to treat PTOA. Furthermore, our results support further exploration of therapeutic strategies that target neuro-metabolic signaling pathways for treating PTOA, particularly in women.

## ACKNOWLEDGEMENTS

The authors gratefully acknowledge Dr. Matthew Silva and Mr. Michael Brodt for teaching us how to conduct the non-invasive compression injury model of PTOA and for sharing the loading fixture designs. We acknowledge Tatsiana Akraiko and Mark Band of the Carver Biotechnology Center, Functional Genomics Lab, University of Illinois for Fluidigm qPCR processing. Finally, we thank Dr. Albert Batushansky for assistance with qPCR data imputation, Ms. Pratibha Dube and Ms. Zainab Sajid for histology assistance, and Dr. Ron June for helpful comments.

This work was supported by the Department of Veterans Affairs (I01BX004666 and 1I01BX004882 Merit Awards to Dr. Griffin) and the Oklahoma Medical Research Foundation. The content is solely the responsibility of the authors and does not necessarily represent the official views of the funding sources, which played no role in the conduct, writing, or submission of the manuscript for publication. The authors have no conflicts of interest to declare.

## AUTHOR CONTRIBUTIONS

Concept and design: RKK, TC, PMD, TMG; acquisition, analysis, and interpretation of data: RKK, PMD, TC, MA, JL, TMG; drafting and critical revision of article: RKK, PMD, TC, MA, JL, TMG; final approval of article: RKK, PMD, TC, MA, JL, TMG.

## DATA SHARING

Data that support the findings of this study are available from the corresponding author upon reasonable request.

## REFERENCES

1. Felson DT, Neogi T. 2018. Emerging Treatment Models in Rheumatology: Challenges for Osteoarthritis Trials. Arthritis & rheumatology (Hoboken, N.J.) 70(8):1175–1181.

2. Scanzello CR. 2017. Chemokines and inflammation in osteoarthritis: Insights from patients and animal models. Journal of Orthopaedic Research 35(4):735–739.

3. Liu-Bryan R, Terkeltaub R. 2014. Emerging regulators of the inflammatory process in osteoarthritis. Nat Rev Rheumatol 11(1):35–44.

4. Christiansen BA, Guilak F, Lockwood KA, et al. 2015. Non-invasive mouse models of post-traumatic osteoarthritis. Osteoarthritis and cartilage 23(10):1627–1638.

5. Anderson DD, Chubinskaya S, Guilak F, et al. 2011. Post-traumatic osteoarthritis: improved understanding and opportunities for early intervention. Journal of Orthopaedic Research 29(6):802–809.

6. Wang Q, Rozelle AL, Lepus CM, et al. 2011. Identification of a central role for complement in osteoarthritis. Nature Medicine 17(12):1674–1679.

7. Griffin TM, Scanzello CR. 2019. Innate inflammation and synovial macrophages in osteoarthritis pathophysiology. Clinical & Experimental Rheumatology 37 Suppl 120(5):57–63.

8. Eming SA, Wynn TA, Martin P. 2017. Inflammation and metabolism in tissue repair and regeneration. Science 356(6342):1026–1030.

9. Courties A, Sellam J, Berenbaum F. 2017. Role of the autonomic nervous system in osteoarthritis. Best Pract Res Clin Rheumatology 31(5):661–675.

10. Sohn R, Rösch G, Junker M, et al. 2021. Adrenergic signalling in osteoarthritis. Cell Signal 82:109948.

11. Rösch G, Bagdadi KE, Muschter D, et al. 2022. Sympathectomy aggravates subchondral bone changes during osteoarthritis progression in mice without affecting cartilage degeneration or synovial inflammation. Osteoarthr Cartilage 30(3):461–474.

12. Lamkin DM, Ho H-Y, Ong TH, et al. 2016. β-Adrenergic-stimulated macrophages: Comprehensive localization in the M1-M2 spectrum. Brain Behav Immun 57:338–346.

13. Ricci MR, Lee M-J, Russell CD, et al. 2005. Isoproterenol decreases leptin release from rat and human adipose tissue through posttranscriptional mechanisms. Am J Physiol-endoc M 288(4):E798–E804.

14. Collins S. 2022. β-Adrenergic Receptors and Adipose Tissue Metabolism: Evolution of an Old Story. Annu Rev Physiol 84(1):1–16.

15. Kosteli A, Sugaru E, Haemmerle G, et al. 2010. Weight loss and lipolysis promote a dynamic immune response in murine adipose tissue. J Clin Investigation 120(10):3466–3479.

16. Valdes AM, Ravipati S, Pousinis P, et al. 2018. Omega-6 oxylipins generated by soluble epoxide hydrolase are associated with knee osteoarthritis. J Lipid Res 59(9):1763–1770.

17. Sebastian A, Hum NR, McCool JL, et al. 2022. Single-cell RNA-Seq reveals changes in immune landscape in post-traumatic osteoarthritis. Front Immunol 13:938075.

18. Christiansen BA, Anderson MJ, Lee CA, et al. 2012. Musculoskeletal changes following non-invasive knee injury using a novel mouse model of post-traumatic osteoarthritis. Osteoarthr Cartilage 20(7):773–782.

19. Seifer DR, Furman BD, Guilak F, et al. 2008. Novel synovial fluid recovery method allows for quantification of a marker of arthritis in mice. Osteoarthritis and Cartilage 16(12):1532–1538.

20. Donovan EL, Lopes EBP, Batushansky A, et al. 2018. Independent effects of dietary fat and sucrose content on chondrocyte metabolism and osteoarthritis pathology in mice. Disease models & mechanisms 11(9):dmm034827.

21. Hahn AK, Batushansky A, Rawle RA, et al. 2021. Effects of long-term exercise and a high-fat diet on synovial fluid metabolomics and joint structural phenotypes in mice: an integrated network analysis. Osteoarthr Cartilage 29(11):1549–1563.

22. Barboza E, Hudson J, Chang W-P, et al. 2017. Profibrotic Infrapatellar Fat Pad Remodeling Without M1 Macrophage Polarization Precedes Knee Osteoarthritis in Mice With Diet-Induced Obesity. Arthritis & Rheumatology 69(6):1221–1232.

23. Man K, Kutyavin VI, Chawla A. 2017. Tissue Immunometabolism: Development, Physiology, and Pathobiology. Cell Metabolism 25(1):11–26.

24. Bossche JV den, O’Neill LA, Menon D. 2017. Macrophage Immunometabolism: Where Are We (Going)? Trends in Immunology 38(6):395–406.

25. June RK, Liu-Bryan R, Long F, Griffin TM. 2016. Emerging role of metabolic signaling in synovial joint remodeling and osteoarthritis. Journal of Orthopaedic Research 34(12):2048– 2058.

26. Berenbaum F, Griffin TM, Liu-Bryan R. 2017. Review: Metabolic Regulation of Inflammation in Osteoarthritis. Arthritis & rheumatology (Hoboken, N.J.) 69(1):9–21.

27. Batushansky A, Zhu S, Komaravolu RK, et al. 2022. Fundamentals of OA. An initiative of Osteoarthritis and Cartilage. Obesity and metabolic factors in OA. Osteoarthr Cartilage 30(4):501–515.

28. Ioan-Facsinay A, Kloppenburg M. 2013. An emerging player in knee osteoarthritis: the infrapatellar fat pad. Arthritis Research & Therapy 15(6):225.

29. Zhou S, Maleitzke T, Geissler S, et al. 2022. Source and hub of inflammation: The infrapatellar fat pad and its interactions with articular tissues during knee osteoarthritis. J Orthop Res 40(7):1492–1504.

30. Hart HF, Culvenor AG, Patterson BE, et al. 2022. Infrapatellar fat pad volume and HoffaLJsynovitis after ACL reconstruction: Association with early osteoarthritis features and pain over 5 years. J Orthop Res 40(1):260–267.

31. Steidle-Kloc E, Culvenor AG, Dörrenberg J, et al. 2018. Relationship Between Knee Pain and Infrapatellar Fat Pad Morphology: A Within-and Between-Person Analysis From the Osteoarthritis Initiative. Arthritis care & research 70(4):550–557.

32. Pan F, Han W, Wang X, et al. 2015. A longitudinal study of the association between infrapatellar fat pad maximal area and changes in knee symptoms and structure in older adults. Ann Rheum Dis 74(10):1818–1824.

33. Han W, Aitken D, Zheng S, et al. 2019. Association Between Quantitatively Measured Infrapatellar Fat Pad High SignalLJIntensity Alteration and Magnetic Resonance Imaging– Assessed Progression of Knee Osteoarthritis. Arthrit Care Res 71(5):638–646.

34. Wang K, Ding C, Hannon MJ, et al. 2018. Signal intensity alteration within infrapatellar fat pad predicts knee replacement within 5 years: data from the Osteoarthritis Initiative. Osteoarthritis and cartilage 26(10):1345–1350.

35. Diepold J, Ruhdorfer A, Dannhauer T, et al. 2015. Sex-differences of the healthy infra-patellar (Hoffa) fat pad in relation to intermuscular and subcutaneous fat content – Data from the Osteoarthritis Initiative. Ann Anat – Anatomischer Anzeiger 200:30–36.

36. Martel-Pelletier J, Tardif G, Pelletier J-P. 2022. An Open Debate on the Morphological Measurement Methodologies of the Infrapatellar Fat Pad to Determine Its Association with the Osteoarthritis Process. Curr Rheumatol Rep 24(3):76–80.

37. Oo WM, Little C, Duong V, Hunter DJ. 2021. The Development of Disease-Modifying Therapies for Osteoarthritis (DMOADs): The Evidence to Date. Drug Des Dev Ther 15:2921– 2945.

38. Fowler-Brown A, Kim DH, Shi L, et al. 2015. The mediating effect of leptin on the relationship between body weight and knee osteoarthritis in older adults. Arthritis & rheumatology (Hoboken, N.J.) 67(1):169–175.

39. Zhu S, Donovan EL, Makosa D, et al. 2022. Sirt3 Promotes Chondrogenesis, Chondrocyte Mitochondrial Respiration and the Development of HighLJFat DietLJInduced Osteoarthritis in Mice. J Bone Miner Res 37(12):2531–2547.

40. Valentine JM, Ahmadian M, Keinan O, et al. 2022. β3-Adrenergic receptor downregulation leads to adipocyte catecholamine resistance in obesity. J Clin Investigation 132(2):e153357.

41. Camell CD, Sander J, Spadaro O, et al. 2017. Inflammasome-driven catecholamine catabolism in macrophages blunts lipolysis during ageing. Nature 550(7674):119–123.

42. Pirzgalska RM, Seixas E, Seidman JS, et al. 2017. Sympathetic neuron–associated macrophages contribute to obesity by importing and metabolizing norepinephrine. Nat Med 23(11):1309–1318.

43. Larabee CM, Neely OC, Domingos AI. 2020. Obesity: a neuroimmunometabolic perspective. Nat Rev Endocrinol 16(1):30–43.

44. Gierman LM, Wopereis S, El B van, et al. 2013. Metabolic profiling reveals differences in concentrations of oxylipins and fatty acids secreted by the infrapatellar fat pad of end-stage osteoarthritis and normal donors. Arthritis & Rheumatism 65(10): 2606–2614.

45. Mustonen A-M, Käkelä R, Finnilä MAJ, et al. 2019. Anterior cruciate ligament transection alters the n-3/n-6 fatty acid balance in the lapine infrapatellar fat pad. Lipids Health Dis 18(1):67.

46. Kosinska MK, Liebisch G, Lochnit G, et al. 2013. A lipidomic study of phospholipid classes and species in human synovial fluid. Arthritis & Rheumatism 65(9):2323–2333.

47. Leimer EM, Pappan KL, Nettles DL, et al. 2017. Lipid profile of human synovial fluid following intra-articular ankle fracture. Journal of Orthopaedic Research 35(3):657–666.

48. Carlson AK, Rawle RA, Wallace CW, et al. 2019. Characterization of synovial fluid metabolomic phenotypes of cartilage morphological changes associated with osteoarthritis. Osteoarthritis and cartilage 27(8):1174–1184.

49. Zhang Y, Vasheghani F, Li Y-H, et al. 2014. Cartilage-specific deletion of mTOR upregulates autophagy and protects mice from osteoarthritis. Ann Rheum Dis 2015;74:1432–1440.

