## Supplemental for "Sex-specific effects of injury and beta-adrenergic activation on metabolic and inflammatory mediators in a murine model of post-traumatic osteoarthritis"

Content:

**Supplemental Methods:** Synovial fluid metabolomic analyses

**Table S1:** 96.96 IFC Custom DELTAgene Panel

**Table S2:** Synovial Fluid Metabolite Statistical Analysis: Effect of Injury

**Figure S1.** Non-invasive single-load knee compression injury model
**Figure S2.** Semi-quantitative scoring criteria to evaluate infrapatellar fat pad (IFP) atrophy and fibrosis following injury **Figure S3:** Treatment-dependent comparison of the effects of injury and biological sex on pro-fibrotic IFP-synovium gene expression
**Figure S4:** Comparison of IFP-synovium *Adrb3* gene expression in males and females across all treatment and injury conditions

**Figure S5:** Injury and sex-dependent comparisons of the effect of isoproterenol treatment on synovial fluid metabolites

**Figure S6:** Injury-dependent effects biological sex on synovial fluid metabolites

**Supplemental Methods**

*Synovial fluid metabolomic analyses*

Synovial fluid was collected from both knees as previously described and stored at -80°C until analysis. Samples were shipped overnight on dry ice to Metabolon (Morrisville, NC, USA) for Global Metabolomic Profiling Analysis. Briefly, samples were prepared using the automated MicroLab STAR® system (Hamilton Company, Reno, NV, USA). Several recovery standards were added prior to the first step in the extraction process for QC purposes. Samples were precipitated with methanol under vigorous shaking for 2 min followed by centrifugation. The resulting extract was divided into five fractions: two for analysis by two separate reverse phase (RP)/UPLC-MS/MS methods with positive ion mode electrospray ionization (ESI), one for analysis by RP/UPLC-MS/MS with negative ion mode ESI, one for analysis by HILIC/UPLC-MS/MS with negative ion mode ESI, and one sample was reserved for backup. Samples were placed briefly on a TurboVap® (Zymark) to remove the organic solvent and stored overnight under nitrogen before analysis. All methods utilized a Waters ACQUITY ultra-performance liquid chromatography (UPLC) and a Thermo Scientific Q-Exactive high resolution/accurate mass spectrometer interfaced with a heated electrospray ionization (HESI-II) source and Orbitrap mass analyzer operated at 35,000 mass resolution. Instrument variability was 5% based on the median relative standard deviation (RSD) for the internal standards added to each sample prior to injection into the mass spectrometers. Overall process variability was 7% based on the median RSD for all endogenous metabolites (i.e., non-instrument standards) present in a pool of human plasma technical replicates. Raw data were extracted, peak-identified and QC processed using Metabolon’s hardware and software. Compound identification was based on comparison to authenticated standard library entries within a narrow retention time/index window, accurate mass to charge ratio (m/z) +/- 10 ppm, and chromatographic data including MS/MS forward and reverse scores between the experimental data and authentic standards. The analysis identified a total of 207 named biochemicals, which were detected in nearly all samples (mean=93.6%, median=100%). Peak area data were normalized to extracted volume and then median scaled. Any missing values (<6.5% of data set) were imputed with sample set minimums on a per biochemical basis.

**Table S1.** 96.96 IFC Array Custom DELTAgene Target List for IFP-Synovium Sample Analyses

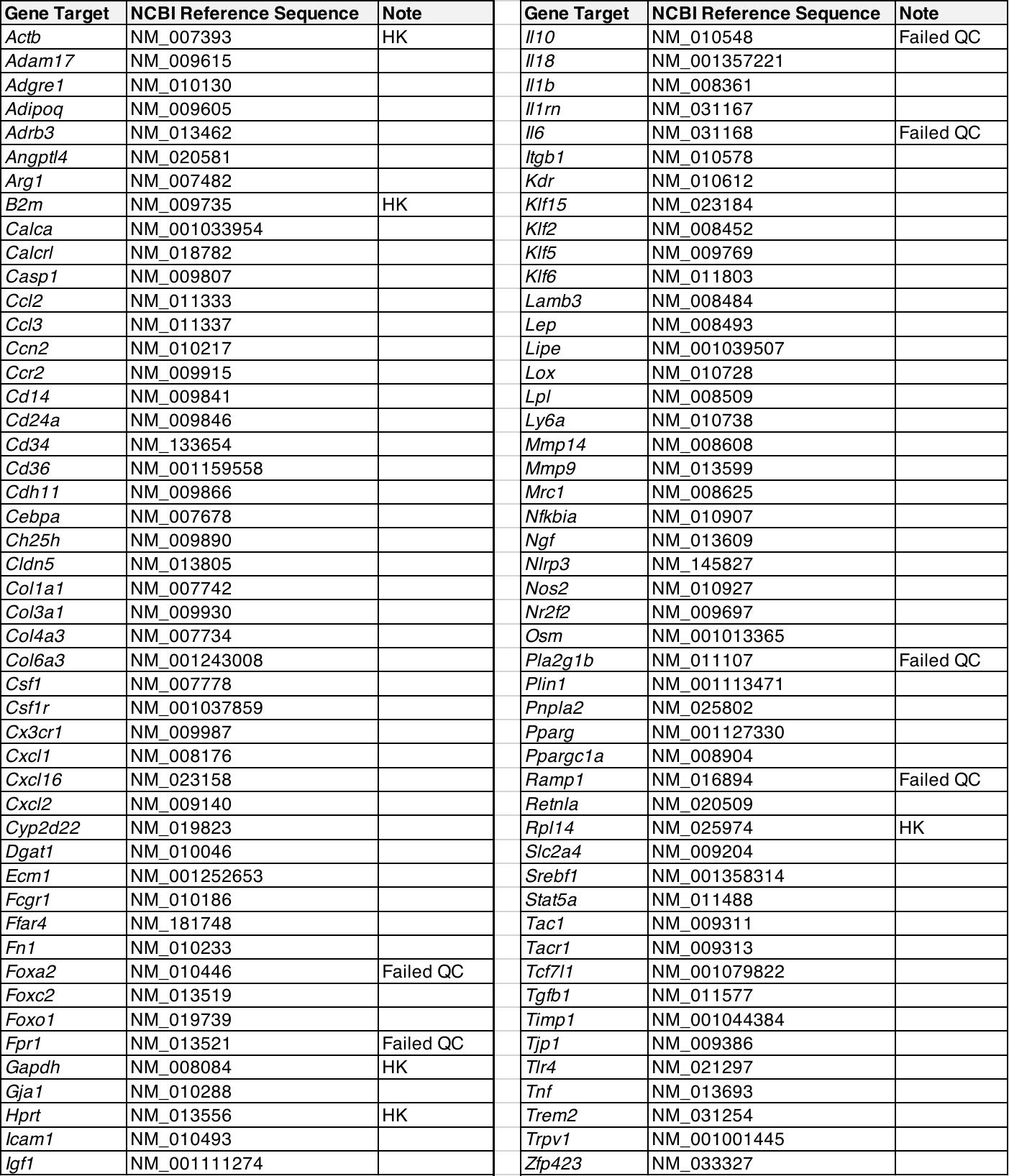

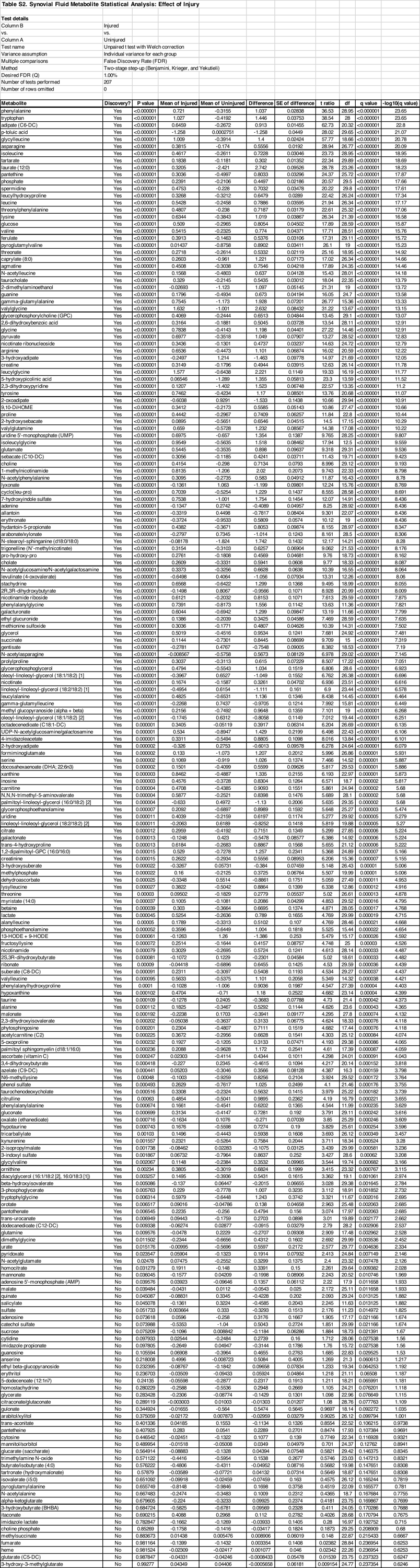

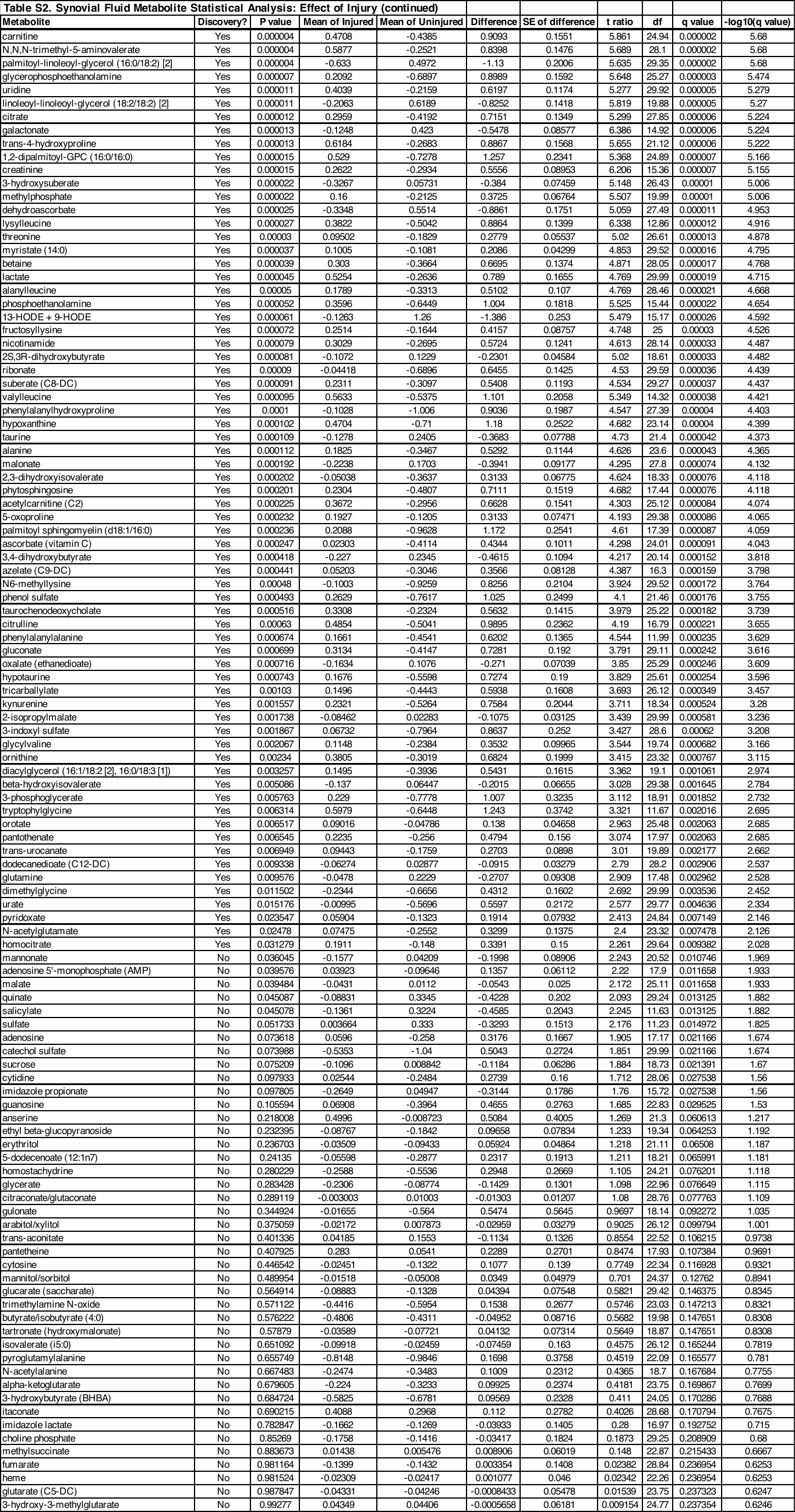

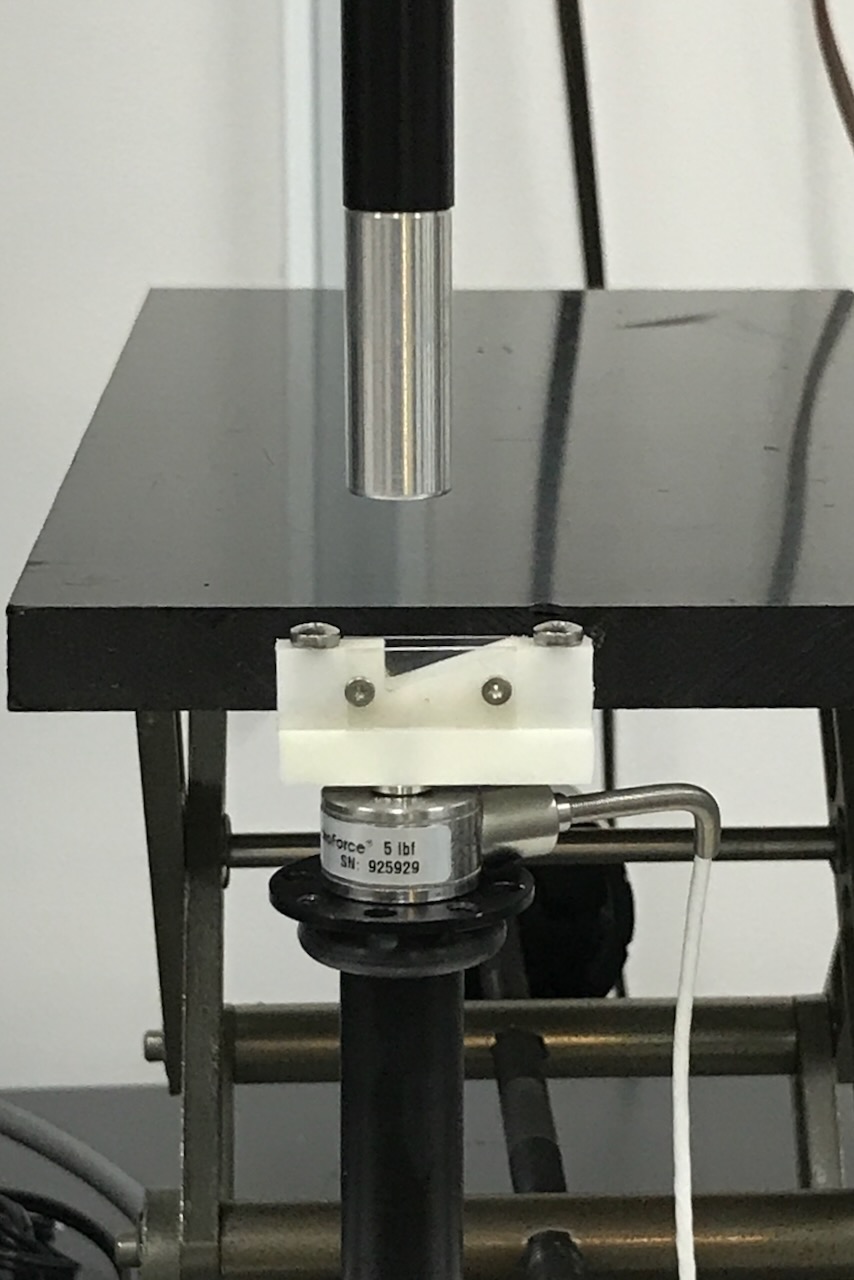

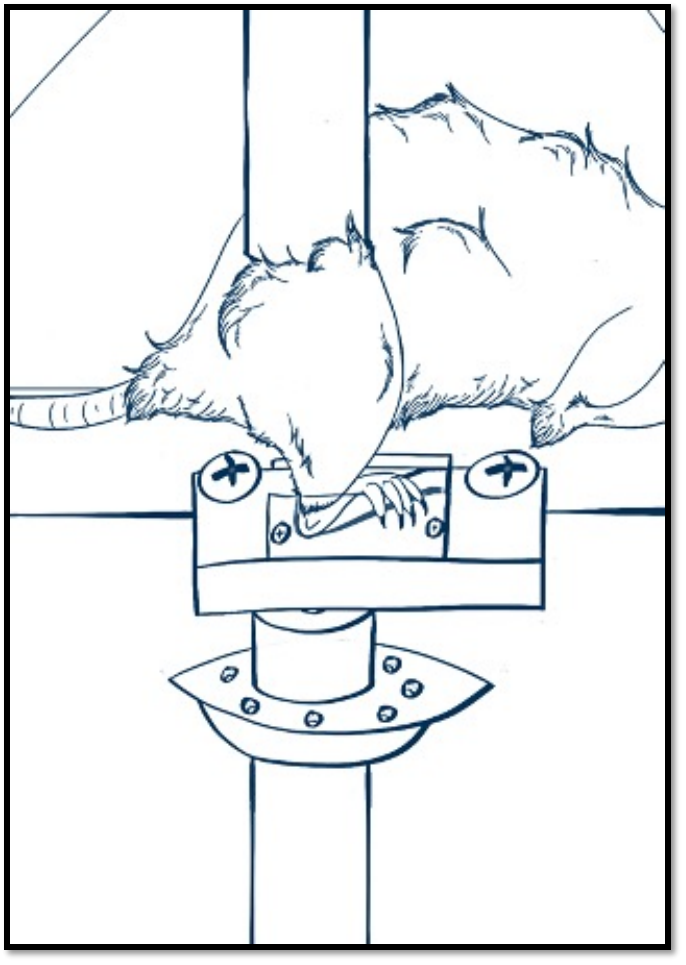

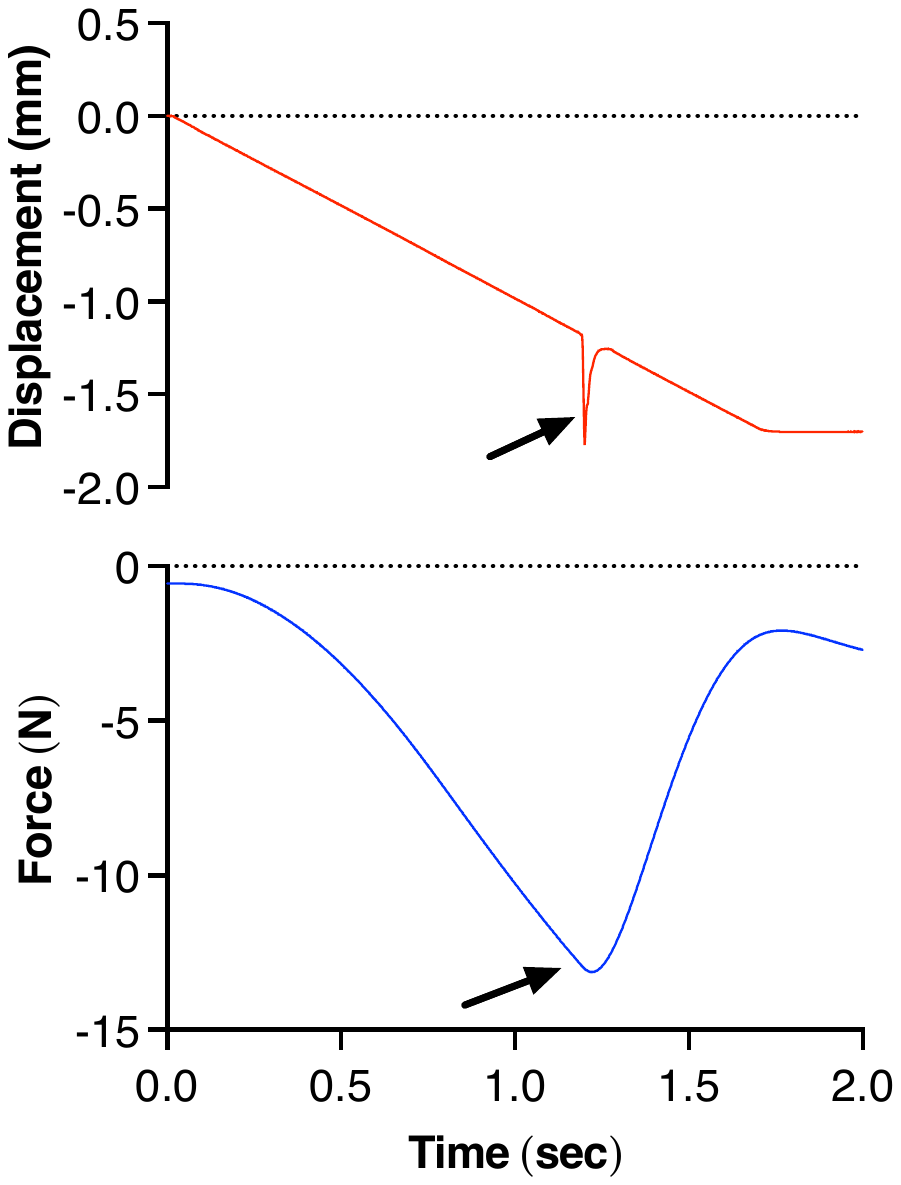

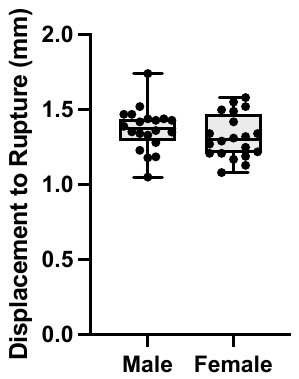

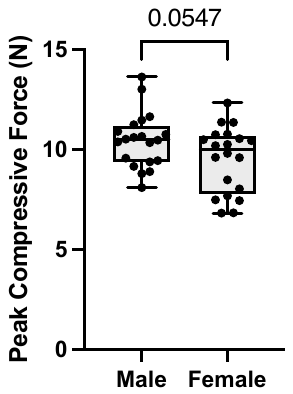

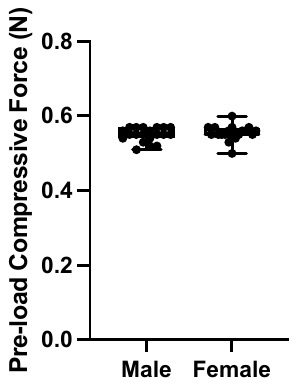

(A)

(B)

(C)

(D)

(E)

(F)

**Figure S1. Non-invasive single-load knee compression injury model.** A. Photo of custom-built foot holding slot secured above a 22N load cell. Custom-built knee holding cup is located directly above foot slot and attached to the actuated upper platen of the electromagnetic materials testing machine. B. Illustration of mouse placement for applying compressive load to the right knee. C. Representative displacement and compression force traces plotted versus time. Arrow indicates the moment of ACL rupture. D. Comparison of pre-load compression force in males and females. E. Comparison of the displacement to generate ACL rupture in males and females. F. Comparison of the peak compressive force resulting from ramp displacement in males and females. Values = mean ± 95% CI. Student’s t-test (two-tailed).

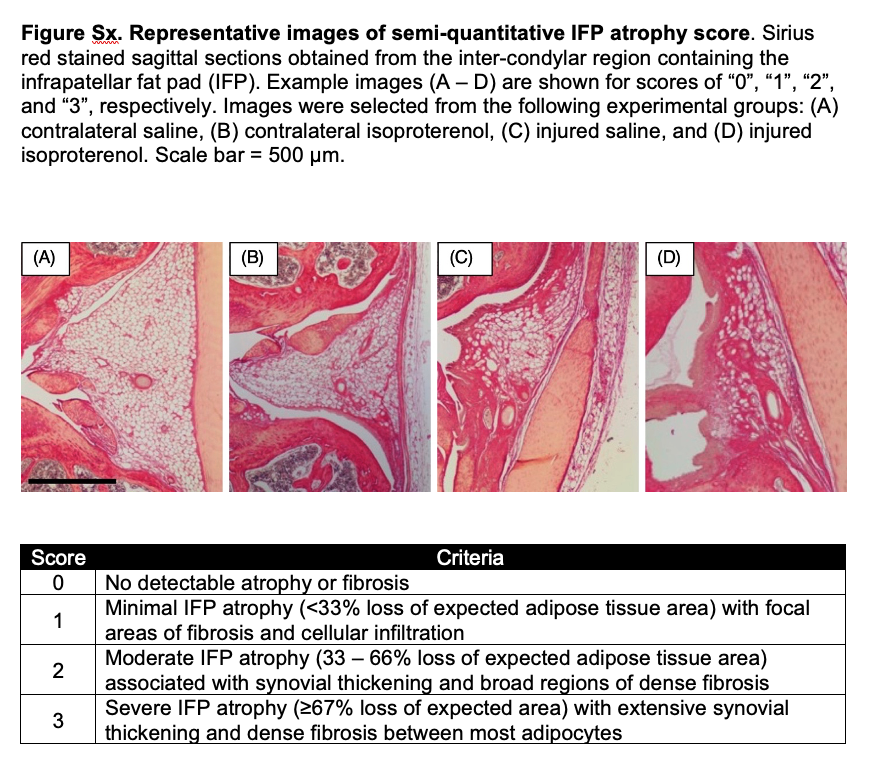

**Figure S2. Semi-quantitative scoring criteria to evaluate infrapatellar fat pad (IFP) atrophy and fibrosis following injury.** Sirius red stained sagittal sections were obtained from the inter-condylar region containing the IFP. Example images (A-D) are shown for scores of “0”, “1”, “2”, and “3”, respectively. Images were selected from the following experimental groups: (A) uninjured (contralateral) saline, (B) uninjured isoproterenol, (C) injured saline, and (D) injured isoproterenol. Scale bar = 500 µm.

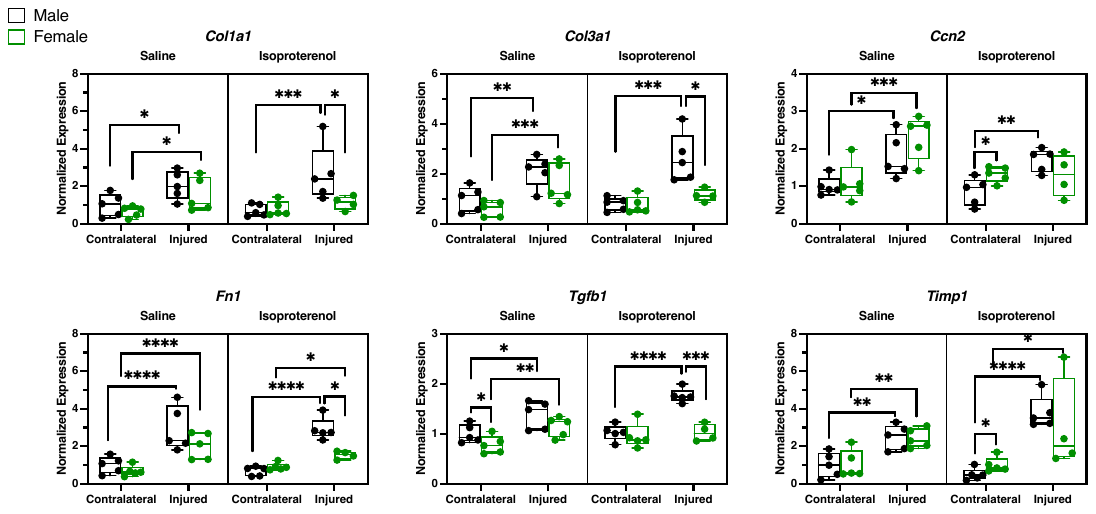

**Figure S3: Treatment-dependent comparison of the effects of injury and biological sex on pro-fibrotic IFP-synovium gene expression.** When the effects of injury and sex were analyzed by 2-factor ANOVA separately for saline and isoproterenol conditions, significant injury-sex interactions (p<0.05) were observed under isoproterenol conditions for half of the fibrosis-associated genes in the target panel (i.e., *Col1a1, Col3a1, Fn1, Tgfb1, Ccn2*, and *Timp1*). These genes were graphed with expression data normalized to the average value of the male contralateral saline treated samples on a per gene basis. Fisher’s LSD post-hoc comparisons are shown for injury and sex comparisons (*p<0.05, **p<0.01, ***p<0.001, ****p<0.0001). Injury increases the expression of these genes under saline conditions in both males and females. However, isoproterenol treatment suppresses the upregulation of these genes following injury in female mice only. Boxes represent the 25th to 75th percentiles, horizontal line indicates the median, and whiskers demonstrate maximum and minimum values. Fisher’s LSD post-hoc paired comparisons shown if p<0.05 for injury and sex comparisons (*p<0.05, **p<0.01, ***p<0.001, ****p<0.0001); n=5 per sex per group, except n=4 for female/injured/isoproterenol.

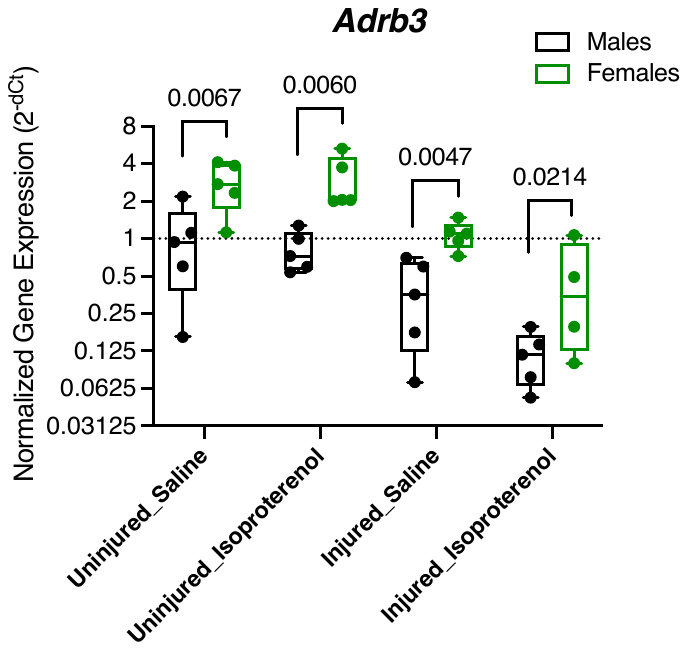

**Figure S4: Comparison of IFP-synovium *Adrb3* gene expression in males and females across all treatment and injury conditions.** Data were normalized to the average value of the male uninjured (i.e., contralateral) saline treated samples, as indicated by the dashed line. *Adrb3*, which is the primary adrenergic receptor isoform expressed in adipocytes, was more highly expressed female versus male samples across all conditions. Injury reduced the expression of *Adrb3*, and isoproterenol treatment also reduced *Adrb3* expression in injured but not uninjured samples. Boxes represent the 25th to 75th percentiles, horizontal line indicates the median, and whiskers demonstrate maximum and minimum values. Fisher’s LSD post-hoc paired comparison p-values shown for condition-specific male - female; n=5 per sex per group, except n=4 for female/injured/isoproterenol.

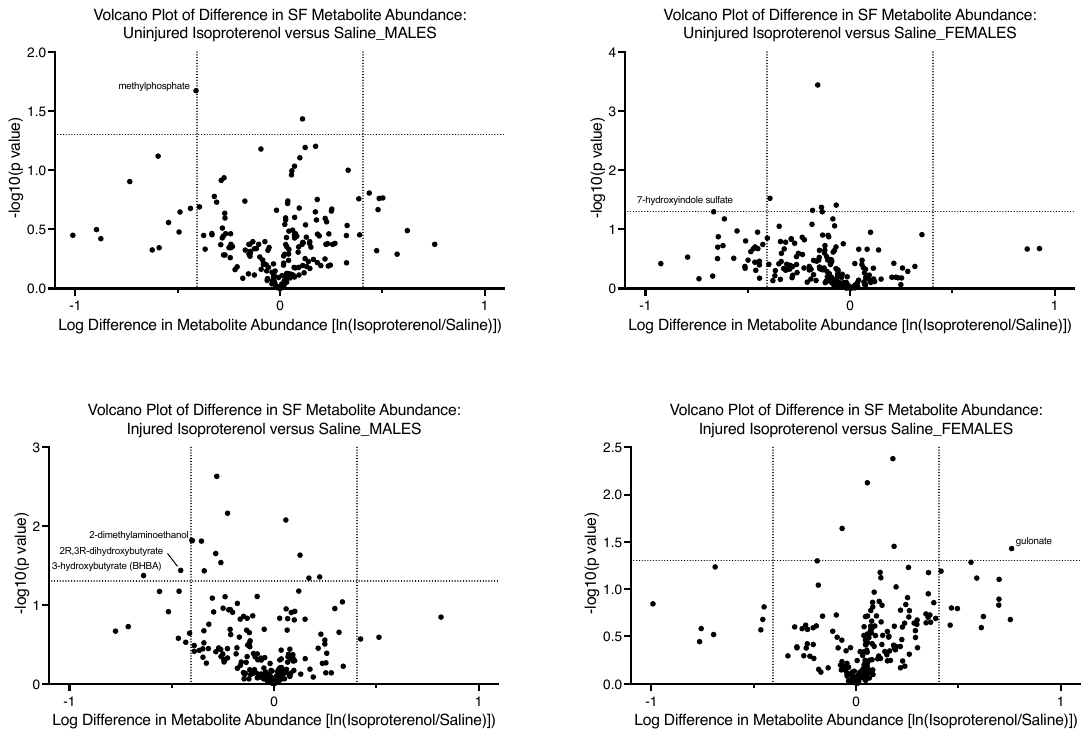

**Figure S5: Injury and sex-dependent comparisons of the effect of isoproterenol treatment on synovial fluid metabolites.** Volcano plot comparisons show minimal effect of isoproterenol treatment on the abundance of synovial fluid metabolites. Vertical dashed lines indicate ±1.5-fold change in metabolite abundance, and the horizontal line indicates a p-value of 0.05 without adjustment for multiple comparisons. For uninjured comparisons, n=3 per sex per treatment group. For injured comparisons, n=5 per sex per treatment group. Metabolites meeting minimal p-value (p<0.05) and magnitude (> ±1.5-fold change) thresholds are labeled.

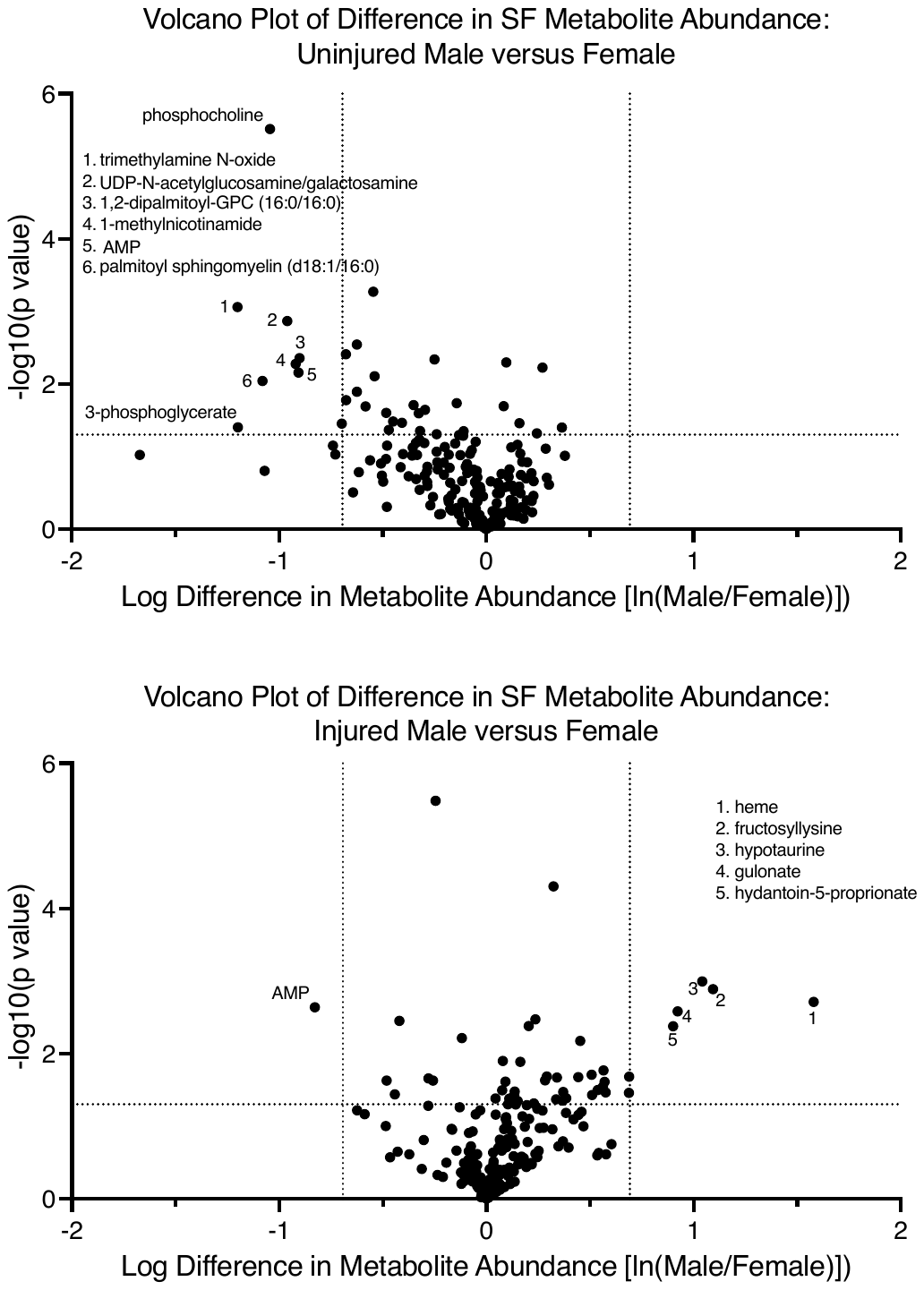

**Figure S6: Injury-dependent effects biological sex on synovial fluid metabolites.** Volcano plot comparisons show modest differences in synovial fluid metabolite abundance between males and females. Comparisons are separated between uninjured and injured conditions. Note, saline and isoproterenol treated samples were pooled due to the negligible effect of isoproterenol (Figure S5) and the benefit of doubling the sample sizes to n=6 per sex for uninjured and n=10 per sex for injured comparisons. Vertical dashed lines indicate ±1.5-fold change in metabolite abundance, and the horizontal line indicates a p-value of 0.05 without adjustment for multiple comparisons. Metabolites meeting minimal p-value (p<0.05) and magnitude (> ±1.5-fold change) thresholds are labeled.
